# Infant Sensory Over-Responsivity: Relations to Early Life Adversity, Emerging Behavioral Problems, and Sensory Functional Connectivity

**DOI:** 10.64898/2026.11.04.764317

**Authors:** Stefanie R. Overschmidt, Hexin Luo, Tara A. Smyser, Aidan Latham, Barbara B. Warner, Cynthia E. Rogers, Deanna M. Barch, Chad M. Sylvester, Christopher D. Smyser, Joan L. Luby, Rebecca F. Schwarzlose

**Author notes:** Correspondence concerning this article should be addressed to Rebecca Schwarzlose, Washington University in St. Louis, 4525 Scott Avenue, St. Louis, MO 63110, United States.

## Abstract

**Background:** Sensory over-responsivity (SOR) in mid- to late childhood is associated with early-life adversity (ELA) and psychopathology, yet the developmental origins and timing of these relationships remain unknown. This work leverages a longitudinal pediatric dataset enriched for ELA to characterize predictors and behavioral problems associated with infant/toddler SOR.

**Methods:** Neonatal fMRI and longitudinal behavioral data were obtained from a sample (N=399) recruited during pregnancy and enriched for social disadvantage (SD). Negative binomial regression models were used to identify pre- and neonatal risk factors for and behavioral correlates of SOR at age two (Y2). Mediation analyses tested whether Y2 SOR mediates relationships between ELA and behavioral problems at age three (Y3). Relationships between neonatal functional connectivity (FC) measures and later SOR were also computed.

**Results:** Two forms of prenatal ELA, SD and maternal psychosocial stress (PSS), positively predicted Y2 SOR, which in turn predicted Y2 and Y3 autism and anxiety problems, greater preschool internalizing symptoms, and greater preschool separation anxiety diagnoses. Positive relationships of prenatal PSS with Y3 autism, anxiety, and internalizing problems were partially mediated by Y2 SOR. Greater neonatal FC between the cortical somatomotor hand network and bilateral thalamus was also associated with SOR at one to two years later.

**Conclusions:** The study results demonstrate the importance of infant/toddler SOR as an early predictor of autism and anxiety behaviors in early childhood and a mediating variable that links prenatal ELA with these behavioral outcomes. Results also implicate heightened neonatal signaling in sensory circuits as an early precursor of SOR.

**Public Significance Statement:** Early life adversity (ELA) is a well-established risk factor for developmental and psychiatric conditions in childhood. Using data from a pediatric sample enriched for ELA, we show that sensory over-responsivity (SOR) in toddlers predicts greater autistic traits and anxiety problems in later childhood and partially mediates relationships between prenatal maternal psychosocial stress and preschool anxiety, autism, and internalizing problems. These results implicate toddler SOR in pathways linking prenatal stress to emerging psychiatric challenges.

---

Sensory over-responsivity (SOR) is an early-emerging behavioral pattern characterized by sensory-elicited distress that can be assessed in infants by one year of age and shows longitudinal stability across development (Ben-Sasson et al., 2010; Carter et al., 2003). Studies of SOR in preschool and school-age children suggest that it is associated with both exposure to adversity and likelihood for neurodevelopmental and psychiatric disorders, particularly autism and anxiety disorders (Baranek et al., 2007; Ben-Sasson et al., 2019; Carpenter et al., 2019; Luo et al., 2026). These associations raise the possibility that early-life adversity may confer risk for behavioral problems, at least in part, by promoting SOR in the infant/toddler period. However, it remains unknown whether SOR in the infant/toddler period mediates relationships between early-life adversity and later behavioral problems. It is also not known whether neural differences in early infancy precede and predict the emergence of SOR in the infant/toddler period. The current study leverages data from a large, longitudinal infant cohort to answer these central questions.

SOR is widely recognized as a feature of autism and was recently added to the diagnostic criteria for autism spectrum disorder (ASD; American Psychiatric Association, 2013). However, SOR is also common in the general population, occurring in approximately 10-20% of typically developing children (Ben-Sasson et al., 2009; Carter et al., 2011; Schwarzlose et al., 2023). Across diverse pediatric samples in middle childhood and adolescence, SOR is consistently and specifically associated with both autistic traits and anxiety symptoms, but not symptoms of other common conditions of childhood (Luo et al., 2026). Longitudinal studies also show that SOR predicts subsequent increases in anxiety symptoms over childhood (Carpenter et al., 2019; Schwarzlose et al., 2023; Tseliou et al., 2025). Like SOR, autism and anxiety typically emerge in early childhood (Baranek et al., 2019; Carpenter et al., 2019; Chen et al., 2022, 2025; Green et al., 2012; McCormick et al., 2016), highlighting the importance of the infant period for understanding the ontogeny and clinical relevance of SOR. To date, few studies have specifically studied behavioral problems associated with SOR in the infant/toddler period; these studies utilized autism-enriched samples to focus on SOR in the context of autism. In particular, a study of toddlers who met criteria for autism (Green et al., 2012) and a prospective study of infants enriched for high familial likelihood of autism (Narvekar et al., 2024) each found that toddler SOR was associated with greater anxiety problems at the preschool period. The results suggest that SOR may precede and predict emerging anxiety, at least among samples enriched for autism or autistic traits. However, it remains unknown whether these effects are specific to autism-enriched samples, or if they generalize to other infant/toddler samples and may be present across the population in early childhood. It is also currently unknown whether infant/toddler SOR mediates relationships between early-life adversity and emerging behavioral problems: a topic that can inform our understanding of the specific pathways by which early adversity promotes lasting neurodevelopmental and psychiatric challenges. The current study leverages data from a longitudinal pediatric sample enriched for social disadvantage (SD), a form of early-life adversity, to address these outstanding questions.

In recent years, there has also been growing interest in using functional MRI to identify the neural correlates of SOR in infancy, which could both inform our understanding of the neurodevelopmental bases of SOR and produce candidate markers for screening. As with studies of infant/toddler SOR and emerging behavioral challenges, work in this space has been primarily conducted in prospective samples enriched for high familial likelihood of autism. Such studies have found differences in functional connectivity (FC) patterns, including that increased FC of thalamic and salience regions with cortical sensory brain regions is specifically associated with high familial likelihood of autism (Nair et al., 2021; Tsang et al., 2024) and, among these infants, specifically with those who exhibit SOR (Wagner et al., 2023). Neuroimaging studies to investigate the neural correlates of SOR in middle childhood and adolescence have also highlighted increased FC of sensory cortical sensory with salience regions and/or the thalamus in boys or across sexes (Cummings et al., 2020; Schwarzlose et al., 2023). Another study comparing FC patterns of children and adults who meet diagnostic criteria for autism and typically developing matched controls also found increased FC between sensory networks and the thalamus (Cerliani et al., 2015). It remains unknown whether similar effects are present in infant/toddler samples that are not enriched for high likelihood of autism, and if so, whether they mediate associations between early-life adversity and SOR. Here, we report results of analyses of resting-state fMRI data collected from the sample in the neonatal period to address these outstanding questions.

It is well-established that children who experience early-life adversity are at heightened risk for developing neurodevelopmental and psychiatric conditions–risk that can compound the deleterious effects of early-life adversity and magnify widespread mental health challenges in adolescence and adulthood (Humphreys & Zeanah, 2014; McLaughlin, et al., 2012; Nelson et al., 2025; Peverill et al., 2021; Reiss, 2013; Vaidya et al., 2024). Given SOR’s links to early adversity and emerging neurodevelopmental and psychiatric conditions, it may represent an intermediate behavioral phenotype, mediating the effects of early adversity on emerging behavioral challenges such as anxiety and autism (Schwarzlose, 2025). Because SOR can be observed in infancy, prior to diagnosis of neurodevelopmental and psychiatric conditions, it may represent an early target for screening and potential intervention to support adaptive sensory processing and mitigate the lasting behavioral effects of early-life adversity. Although early-life adversity is a well-known risk factor for both SOR and lifelong mental health challenges, no prior study has investigated the relationships among early adversity, SOR, FC patterns, and emerging behavioral challenges. Using a longitudinal sample enriched for SD, we aim to address this knowledge gap and characterize the specific behavioral and biological pathways linking early-life adversity and SOR to behavioral outcomes.

## Methods

### Participants

Data were obtained from the longitudinal eLABE (Early Life Adversity Biological Embedding and Risk for Developmental Precursors of Mental Disorders) Study at Washington University in St. Louis, which collects behavioral data from pregnant mothers beginning prenatally and both behavioral and neuroimaging data from their children beginning after birth. Women who participated in a study of preterm birth with the Prematurity Research Center at Washington University in St. Louis were recruited during pregnancy, primarily during the third trimester, from two clinics, including one for Medicaid-insured women specifically. The total participant pool was over-sampled for SD. Exclusion criteria included pregnancy complications, fetal congenital anomalies or infections, multiple gestations, and prenatal alcohol or substance use, except for tobacco and/or marijuana (Luby et al., 2023). This study was approved by the Washington University School of Medicine Institutional Review Board. Analyses for the current study were limited to infants born at gestational age at or above 37 weeks, omitting 56 infants from subsequent analysis due to a relatively low number of preterm infants in the sample, such that preterm infants would only make up 14-39 additional infants in any given analysis. Given the underrepresentation of preterm infants, we opted not to include this additional source of variation in preterm birth due to limited power (see Figure S1a). The present sample was comprised of 56.36% male and 43.64% female infants. Mean income-to-needs ratio was 2.96, and median income-to-needs ratio was 1.39. Racial and ethnic makeup were as follows: 39.41% white, 59.75% African American, 1.27% Chinese, 0.42% other Pacific Islander, 0.42% other (1.27% of participants reported multiple races); 96.61% non-Hispanic, 2.54% Hispanic (0.85% of respondents declined to answer). Supplemental analyses of infants born at gestational age at or above 30 weeks were also performed and are reported in the supplement (n = 394). Timepoints used for analyses in the current study included prenatal, neonatal, year 1 (Y1), year 2 (Y2), year 3 (Y3), and preschool/school-age collected between four and six years of age. Sample sizes for specific analyses vary by analysis based on data availability for individual measures, as reported in tables and supplement (see Table S1). Because this is a secondary analysis of an existing dataset, we do not have control over sample size and, therefore, the present sample could be underpowered to detect small-to-moderate effects. This paper interprets positive results, and the authors cannot speak to whether failure to find significant effects are the result of sample-size limitations.

### Adversity and Behavioral Measures

#### Adversity measures

Latent factors of two forms of early-life adversity (prenatal SD and prenatal maternal psychosocial stress) are computed from questionnaire and demographic data, as described elsewhere (Luby et al., 2023; Supplement).

#### Child behavioral measures

Behavioral measures were based on the Infant-Toddler Social and Emotional Assessment (ITSEA) administered at years one and two (Carter et al., 2003), Child Behavior Checklist (CBCL) 1.5-5 years administered at years two and three (Achenbach & Rescorla, 2000), and second-edition Preschool Age Psychiatric Assessment (PAPA) administered between years four and six (Egger et al., 2020). The Sensory Sensitivity Subscale of the ITSEA is used as a measure of SOR in the current analyses (see Table S2 for subscale items). Analyses focused on Y2 SOR rather than Y1 SOR because of prior work showing stability in sensory symptoms from age 2 through middle childhood in children with autism (McCormick et al., 2016). Internalizing and externalizing domain raw scores were used to broadly assess emerging behavioral problems related to internalizing and externalizing disorder in childhood. Raw scores from five DSM-oriented scales measuring affective problems, anxiety problems, pervasive developmental problems, attention deficit/hyperactivity problems, and oppositional defiant problems were used to assess emerging disorder-specific behavioral problems (see Table S3 for subscale items). The pervasive developmental problems scale, also known as the autism spectrum problems scale, has been validated as an effective screening tool for autism spectrum disorder (Levy et al., 2019; Rescorla et al., 2019). Diagnostic measures were obtained from the second-edition Preschool Age Psychiatric Assessment (PAPA), a structured parent-report diagnostic interview (Egger et al., 2020). Both number of symptoms and diagnostic status were used to assess psychiatric problems in both the internalizing and externalizing domains, as well as those related to Generalized Anxiety Disorder, Social Phobia, and Separation Anxiety Disorder. See Supplement for additional details on behavioral measures.

#### Neuroimaging measures

Neuroimaging measures were collected with MRI in the first weeks of life on a 3T Prisma scanner with a 64-channel Siemens head coil. Infants were scanned without sedation during natural sleep. MRI images collected during scans included T2-weighted anatomical images (voxel size 0.8 mm isotropic, echo time [TE] = 563 ms, repetition time [TR] = 3.2 sec or 4.5 sec) and gradient-recalled echo-planar sequences for BOLD fMRI (TR = 800 ms, TE = 37, multi-band factor = 8, voxel size 2.0 mm isotropic). The preprocessing pipeline has been described elsewhere (Tooley et al., 2025) and included motion correction, distortion correction, intensity normalization, and linear registration of functional images to individual T2 images and then cohort group-average T2 images in 711-2N Talairach atlas space. The MCRIB surface-based neonatal parcellation was used to generate cortical surfaces for individual participants (Alexander et al., 2017). Functional data were frame-censored with a threshold of framewise displacement > 0.25 mm and a minimum of 3 consecutive uncensored frames required for data retention, bandpass filtered between 0.005 Hz and 0.1 Hz, demeaned and detrended. Spherical registration was used to align fMRI data into the Freesurfer 32k surface space across participants. Functional data were parcellated based on the Gordon-Laumann parcellation and network communities derived from adult participants (Gordon et al., 2016). A subset of these network parcels, including many parcels in somatosensory and auditory networks, have been shown to perform well in characterizing infant FC patterns (Tu et al., 2025). Summary (mean) FC measures were computed in a pairwise fashion between network communities and bilateral subcortical structures.

Data and study materials are not publicly available but may be available upon request. Analysis code used for this paper is available upon request. This work was not preregistered.

### Analyses

#### Behavioral Analyses

Given non-normal and zero-skewed distributions of ITSEA and CBCL subscores in the sample, negative binomial models were used for all behavioral analyses (see supplement for more information). Such analyses were conducted in R (v. 4.4.3) using the glm.nb function from the MASS package (Venables & Ripley, 2002). All measures were winsorized using the 90% winsorization method prior to analysis to mitigate the impact of outliers on analysis results. Winsorization was performed in R (v. 4.4.3), using both base R and the filter function from the dplyr package (Wickham et al., 2025).

Negative binomial models of Y2 SOR were computed with candidate risk factors (i.e., SD, PSS, birthweight, gestational age, child sex, and Y1 SOR) as predictors in independent bivariate analyses. For those relationships that were found to be significant (i.e, Y2 SOR with SD, PSS, and Y1 SOR), Y2 age and, separately, sex were included as covariates in subsequent analyses. Associations between Y2 SOR and concurrent (Y2) internalizing problems, externalizing problems, and DSM-oriented problem scales were also conducted in models of Y2 SOR predicting Y2 behavioral problems. Correction for multiple comparisons across internalizing and externalizing problem domains and, separately, across the five DSM-oriented problem scales was performed using the Bonferroni method with the step-up correction (Hochberg, 1988; Menyhart et al., 2021). Significant relationships were subjected to additional post-hoc testing in models with SD (given sample enrichment for disadvantage), Y2 age, and sex separately included as covariates (see Table S4 for full list of regression analyses performed).

The predictive role of Y2 SOR in the emergence of Y3 symptoms was assessed with Y3 CBCL DSM-oriented scales and Y3 CBCL internalizing and externalizing scales as outcome variables. Correction for multiple comparisons was applied across the internalizing and externalizing problem domains and, separately, across the five DSM-oriented problem scales using the Bonferroni method with the step-up correction (Hochberg, 1988). For those relationships that were found to be significant, follow-up analyses with models that included SD, Y2 age, Y3 age, and sex were computed to test whether effects were robust to inclusion of these covariates.

The predictive role of Y2 SOR in the symptom profile and diagnosis of psychiatric disorders at preschool-age was assessed using the number of symptoms and diagnostic status from the PAPA, administered between four and six years of age. Number of symptoms and, separately, diagnostic status of any internalizing disorder, any externalizing disorder, Generalized Anxiety Disorder, Social Phobia, and Separation Anxiety Disorder were outcome variables. For effects found to be significant, follow-up analysis with a model including age at PAPA administration was computed to test robustness of this effect to covariate selection.

Mediation models were additionally computed in R using negative binomial analyses to assess the mediating role of Y2 SOR in mediating the relationships between prenatal SD and, separately, prenatal PSS, with Y3 symptoms significantly predicted by Y2 SOR in previous analyses (i.e., anxiety, autism spectrum, internalizing, and externalizing symptoms).

#### Functional Connectivity Analyses

General linear models of neonatal pairwise network-level FC were computed in base R to assess the relationship between SOR and neonatal FC measures in the following *a priori* network-level pairs: caudate-cinguloparietal network, thalamus-auditory cortex, caudate-auditory cortex, thalamus-somatomotor hand network, caudate-somatomotor hand network, somatomotor hand network-somatomotor hand network, somatomotor mouth network-somatomotor hand network, salience network-somatomotor hand network, ventral attention network-ventral attention network. Network-level pairs were selected for *a priori* testing based on prior findings that they are related to SOR in preadolescents (Schwarzlose et al., 2023). Given the strong relationship between Y1 and Y2 SOR scores, a composite measure of SOR scores from these two timepoints was used to maximize available sample size (see supplement for additional information). Correction for multiple comparisons was performed using the Bonferroni method with the step-up correction (Hochberg, 1988). For those relationships found to be significant in bivariate models, separate post-hoc models containing SD, PSS, or number of frames retained as covariates were additionally performed.

## Results

### Predictors of SOR at two years of age

Prenatal SD and PSS were each significantly associated with Y2 SOR (βs=.30; ps≤.01; Table 1, Figure 1; see Figure S2 for correlations across all major variables). By contrast, birthweight, gestational age at birth, and child sex were not significant predictors of Y2 SOR (Table 1, Figure S1, Supplement for follow-up analyses). Analyses indicate continuity of SOR across infant timepoints, such that Y1 SOR significantly predicted Y2 SOR (β=1.41, p<.001; Figure S3). Post-hoc sensitivity analyses and follow-up analyses including preterm infants (GA≥30 weeks at birth) from the sample yielded similar patterns of results, suggesting the observed results are largely robust to covariate and sample selection (Table S5, Table S6, Supplement).

**Figure 1.**
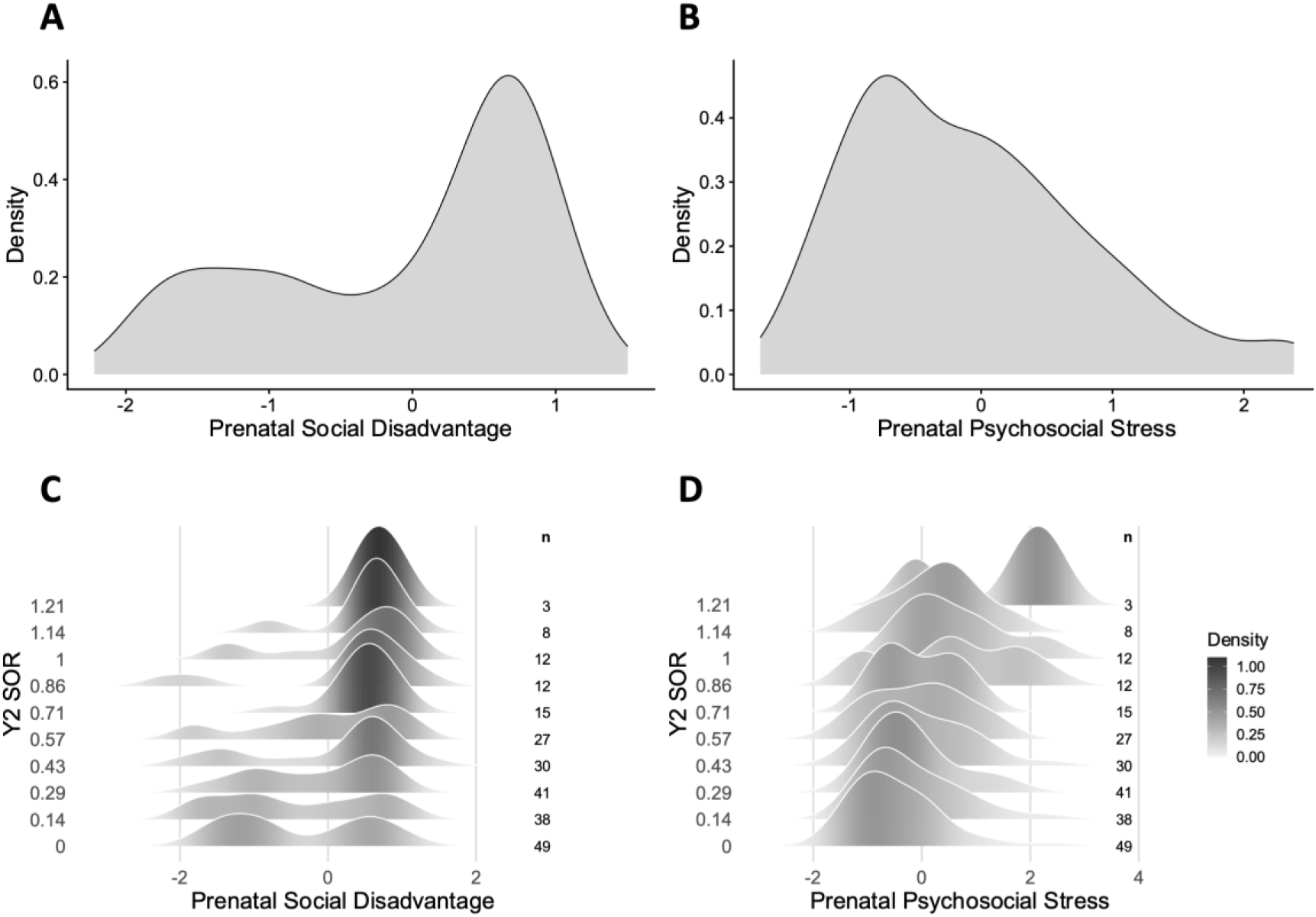
Density of Early Life Adversity Measures and Their Relation to SOR at Two Years of Age Note: (Top) Density plots of A) winsorized prenatal social disadvantage and B) winsorized prenatal maternal psychosocial stress in the sample. Higher latent factor values indicate greater adversity. (Bottom) Density plots of C) prenatal social disadvantage and D) prenatal maternal psychosocial stress stratified by winsorized Y2 SOR raw scores on y-axis. Density is indexed by height and shading, with denser regions of the distribution shown in darker shades.

**Table 1.**
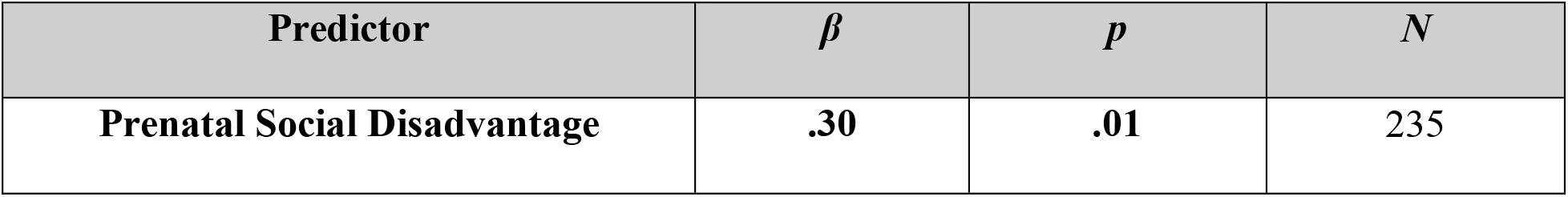
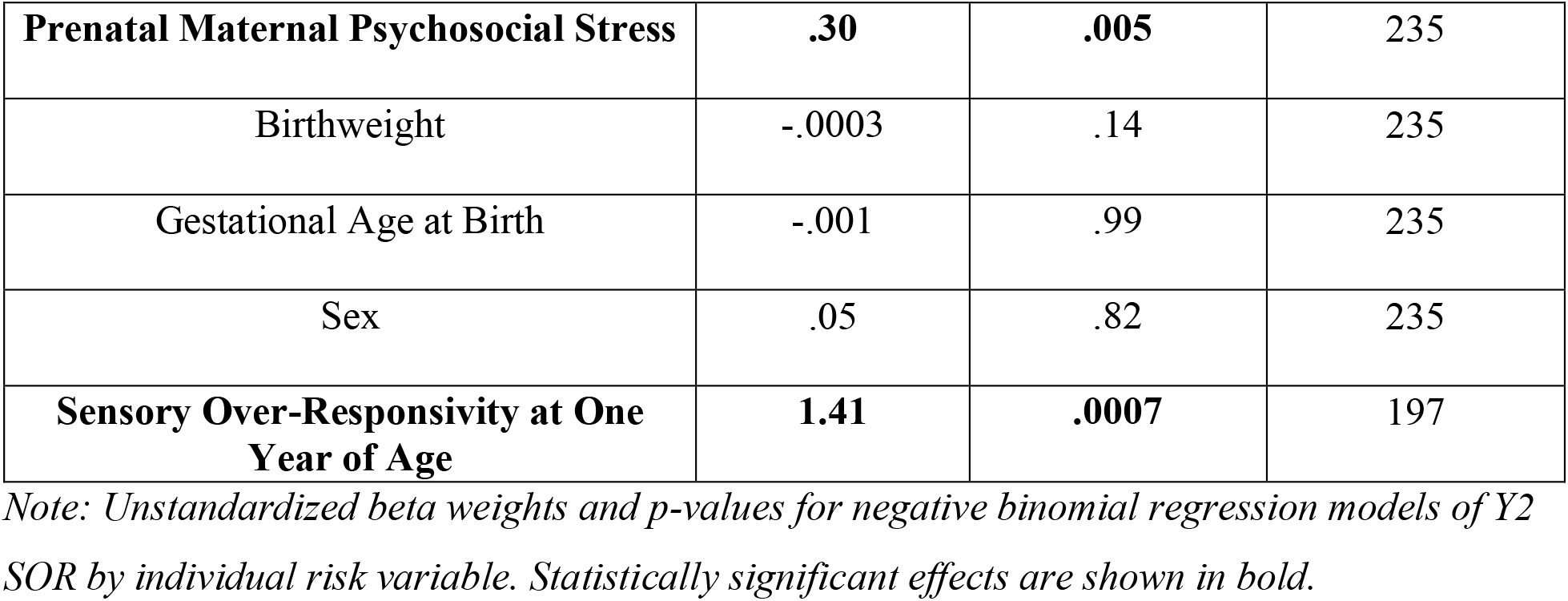
Early Life Adversity and Demographic Variables in Relation to SOR at Two Years of Age.

### Relations between SOR and emerging behavioral problems

#### Concurrent behavioral problems correlated with SOR at two years of age

Subsequent analyses examined associations between SOR and behaviors associated with emerging psychiatric problems (Figure S4). Y2 SOR was significantly associated with both concurrent internalizing problems (β=1.06, p<.000001) and externalizing problems (β=.53, p<.05); both effects remained significant after correction for multiple comparisons and in follow-up sensitivity analyses, demonstrating that effects were robust to covariate selection (Table 2, Table S7, Supplement).

**Table 2.**
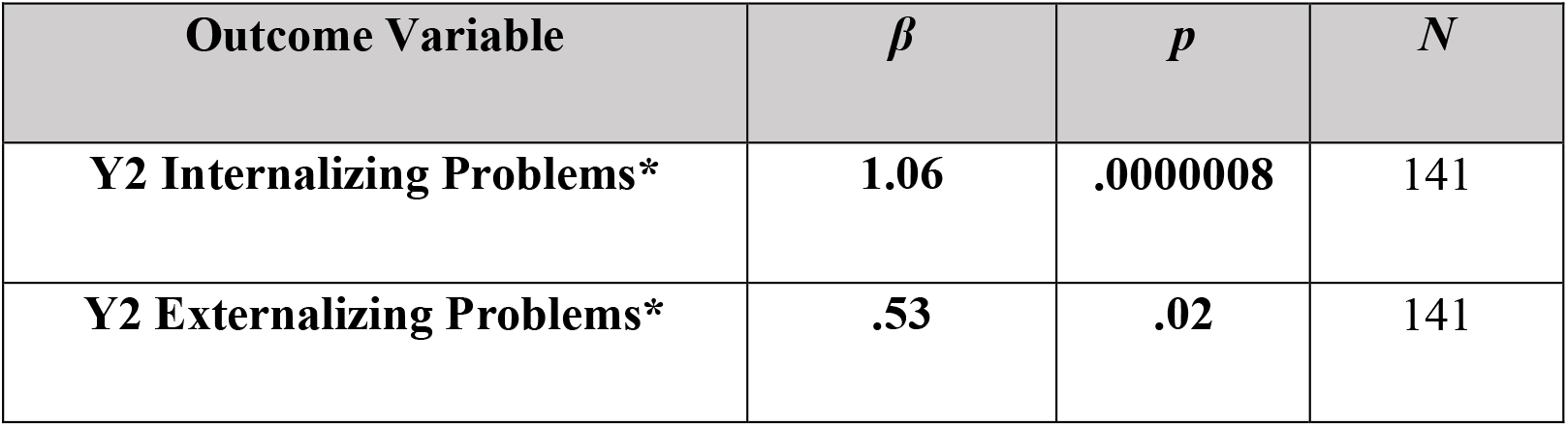
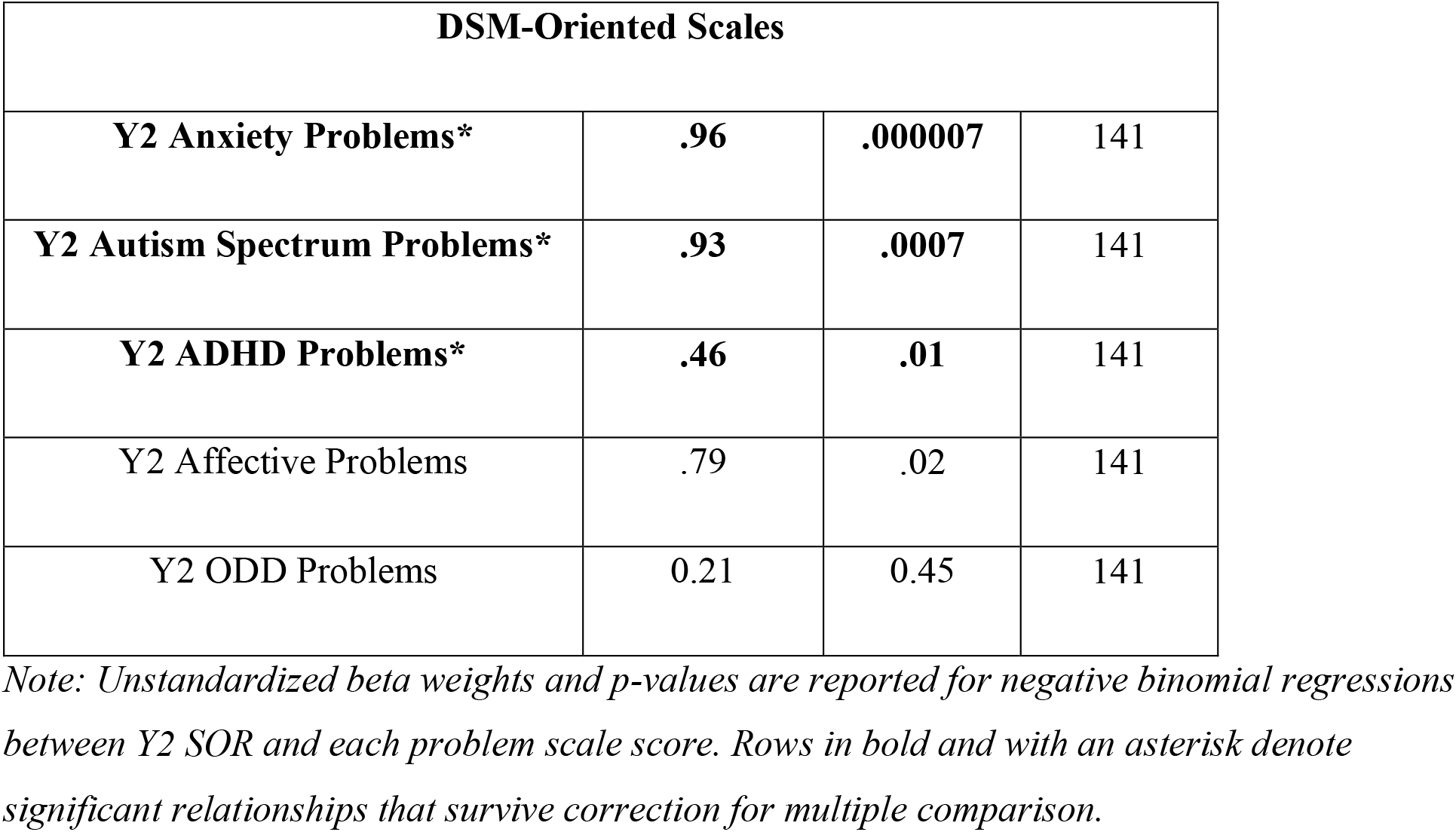
SOR at Two Years Is Associated with Specific Concurrent Behavioral Problems.

| **Outcome Variable** | ***β*** | ***p*** | ***N*** |
|---|---|---|---|
| **Y2 Internalizing Problems\*** | **1.06** | **.0000008** | 141 |
| **Y2 Externalizing Problems\*** | **.53** | **.02** | 141 |
| **DSM-Oriented Scales** |  |  |  |
| **Y2 Anxiety Problems\*** | **.96** | **.000007** | 141 |
| **Y2 Autism Spectrum Problems\*** | **.93** | **.0007** | 141 |
| **Y2 ADHD Problems\*** | **.46** | **.01** | 141 |
| Y2 Affective Problems | .79 | .02 | 141 |
| Y2 ODD Problems | 0.21 | 0.45 | 141 |

*Note: Unstandardized beta weights and p-values are reported for negative binomial regressions between Y2 SOR and each problem scale score. Rows in bold and with an asterisk denote significant relationships that survive correction for multiple comparison.*

Among DSM-oriented subscale scores, Y2 SOR was specifically related to greater concurrent anxiety problems (β=.96, p<.00001), autism spectrum problems (β=.93, p<.001), and ADHD problems (β=.46, p<.05) but not other subscales after correction for multiple comparisons (See Table 2). Post-hoc sensitivity analyses confirmed that the observed effects were robust to covariate selection; see Table S7 and Supplement. Results from follow-up analysis of the inclusive sample (GA≥30 weeks) were the same, except that ADHD problems were not significantly related to SOR (Tables S8 and S9).

#### SOR at age two predicts behavioral problems at age three

Y2 SOR significantly predicted Y3 internalizing, externalizing, anxiety, autism spectrum, and affective problems after correction for multiple comparisons (Table 3). In follow-up analyses that included Y2 behavioral scores as covariates, Y2 SOR remained significantly associated with Y3 internalizing, autism spectrum, and affective problems, indicating that *Y2 SOR predicts increases in these behavioral problems* between two and three years of age (Table 3). These effects proved robust to covariate selection in post-hoc sensitivity analyses (Tables S10 and S11). The strongest effects (Y2 SOR predicting greater Y3 internalizing, anxiety, and autism spectrum problems) were also significant in follow-up analyses including preterm infants (Tables S12 and S13).

**Table 3.** SOR at Age Two Predicts Specific Behavioral Problems at Age Three.

|  | **Bivariate Model** |  |  | **Y2 Sx included as covariate** |  |  |
|---|---|---|---|---|---|---|
| Outcome Variable | ***β*** | ***p*** | *N* | ***β*** | ***p*** | *N* |
| **Y3 Internalizing Problems\*** | **.87** | **.0000007** | 164 | **.79** | **.001** | 96 |
| **Y3 Externalizing Problems\*** | **.41** | **.02** | 164 | .39 | .08 | 96 |
| **DSM-Oriented Scales** |  |  |  |  |  |  |
| **Y3 Anxiety Problems\*** | **.81** | **.00007** | 164 | .43 | .08 | 96 |
| **Y3 Autism Spectrum Problems\*** | **.94** | **.000003** | 164 | **.87** | **.003** | 96 |
| Y3 ADHD Problems | .35 | .02 | 164 | .36 | .07 | 96 |
| **Y3 Affective Problems\*** | **.58** | **.01** | 164 | **.66** | **.02** | 96 |
| Y3 ODD Problems | .35 | .09 | 164 | - | - | - |

*Note: Unstandardized beta weights and p-values are reported for negative binomial regressions between Y2 SOR and each Y3 problem scale score. Rows in bold and with asterisks denote significant relationships that survived multiple comparison correction.*

#### SOR at age 2 predicts preschool symptoms in a clinical interview

Subsequent analyses examined whether Y2 SOR predicted psychiatric symptoms and diagnoses obtained via the PAPA clinical interview in the preschool period (i.e., between four and six years of age). Autism diagnoses were not available at time of writing. Y2 SOR was significantly associated with greater preschool internalizing symptoms (β=.40, p<.05) and separation anxiety symptoms (β=.71, p<.01) after correction for multiple comparison. Positive relations were also observed between Y2 SOR and preschool social phobia and generalized anxiety symptoms, however these effects did not survive correction for multiple comparisons (Table 4). Post-hoc sensitivity analyses confirmed that results were robust to differences in participant age at PAPA administration (Table S14). In follow-up analyses that included children born preterm, Y2 SOR remained a significant predictor of preschool separation anxiety symptoms (Table S15, Supplement).

**Table 4.** Y2 SOR Predicts Internalizing and Specific Anxiety Symptoms in the Preschool Period.

|  | **Bivariate Model** |  |  |
|---|---|---|---|
| **Outcome Variable** | ***β*** | ***p*** | ***N*** |
| **Number of Internalizing Symptoms\*** | **.40** | **.02** | 180 |
| Number of Externalizing Symptoms | .21 | .44 | 180 |
| **Number of Separation Anxiety Symptoms\*** | **.71** | **.009** | 180 |
| Number of Social Phobia Symptoms | 1.37 | .03 | 180 |
| Number of Generalized Anxiety Symptoms | .38 | .03 | 180 |

*Note: Unstandardized beta weights and p-values are reported for negative binomial regressions between Y2 SOR and each symptom count. Rows in bold and with asterisks denote significant relationships that survived multiple comparison correction.*

Y2 SOR also significantly predicted preschool-age separation anxiety diagnoses after correction for multiple comparisons (β=1.45, p<.05). A second, positive relationship between Y2 SOR and preschool-age social phobia diagnoses did not survive multiple-comparison correction (Table 5). The relationship between Y2 SOR and preschool separation anxiety diagnosis was robust in post-hoc sensitivity analyses (Table S16) but failed to reach significance in analyses that included children born preterm (Table S17).

**Table 5.**
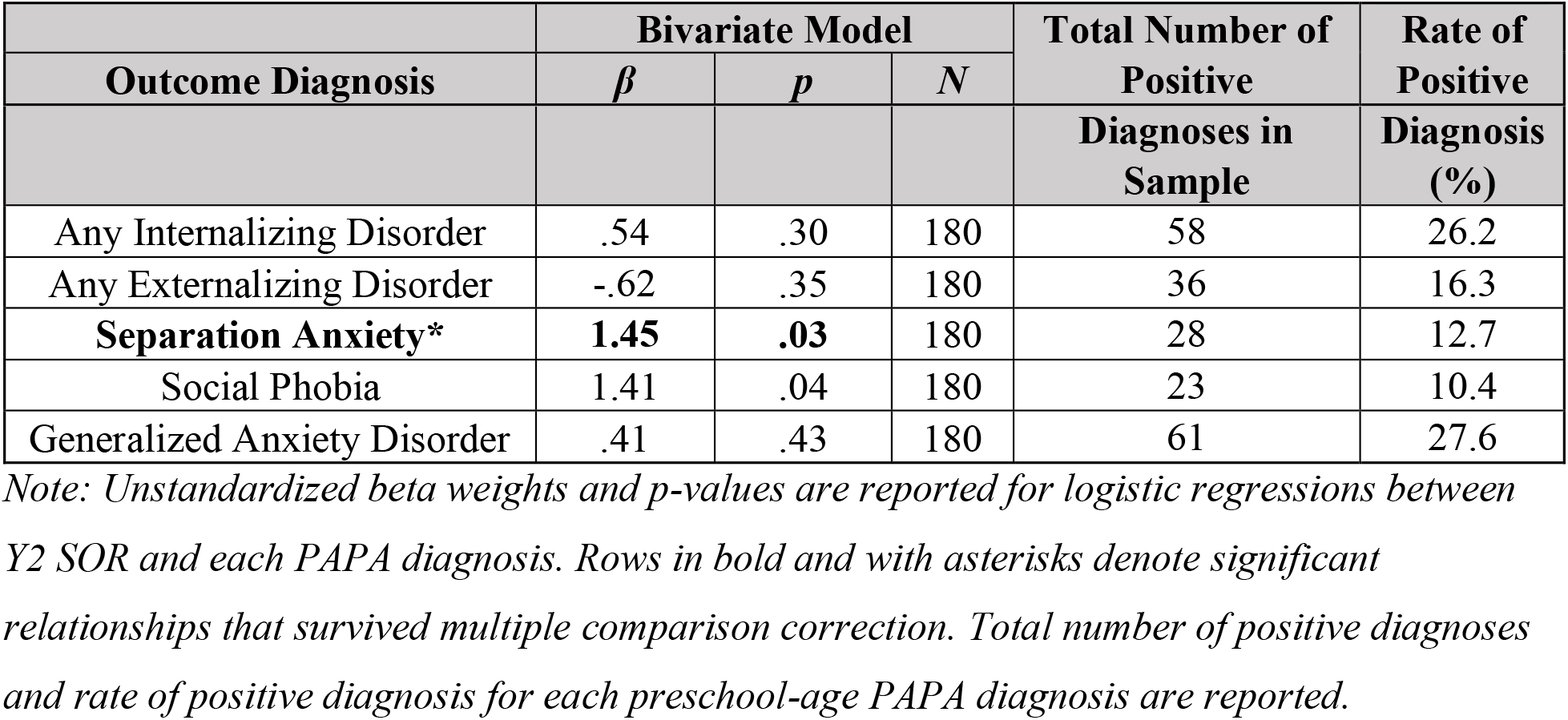
SOR at Age Two Predicts Separation Anxiety Diagnosis in the Preschool Period.

| **Outcome Diagnosis** | **Bivariate Model** |  |  | **Total Number of Positive** | **Rate of Positive** |
|---|---|---|---|---|---|
|  | ***β*** | ***p*** | ***N*** |  |  |
|  |  |  |  | **Diagnoses in Sample** | **Diagnosis (%)** |
| Any Internalizing Disorder | .54 | .30 | 180 | 58 | 26.2 |
| Any Externalizing Disorder | -.62 | .35 | 180 | 36 | 16.3 |
| **Separation Anxiety\*** | **1.45** | **.03** | 180 | 28 | 12.7 |
| Social Phobia | 1.41 | .04 | 180 | 23 | 10.4 |
| Generalized Anxiety Disorder | .41 | .43 | 180 | 61 | 27.6 |

*Note: Unstandardized beta weights and p-values are reported for logistic regressions between Y2 SOR and each PAPA diagnosis. Rows in bold and with asterisks denote significant relationships that survived multiple comparison correction. Total number of positive diagnoses and rate of positive diagnosis for each preschool-age PAPA diagnosis are reported.*

#### SOR as a mediator of ELA-symptom associations

Next, we tested whether SOR mediates relationships between prenatal adversity factors and Y3 autism spectrum and anxiety problems in this sample. Whereas PSS positively predicted Y3 autism spectrum, anxiety, internalizing and externalizing problems (βs≥.21, ps<.001), SD did not significantly predict any of these Y3 measures (ps >.05). Therefore, analyses tested whether Y2 SOR mediated the relationships between prenatal PSS and these behavioral problems at Y3.

#### Prenatal maternal psychosocial stress

Y2 SOR partially mediated the relationship between PSS and each of the following measures: Y3 anxiety, autism spectrum, and internalizing problems (Figure 2, Figure S5). In contrast, Y2 SOR did not significantly mediate the relationship of PSS with Y3 externalizing problems. Follow-up analyses including children born preterm yielded the same pattern of results (Figure S6, Figure S7, Supplement).

**Figure 2.**
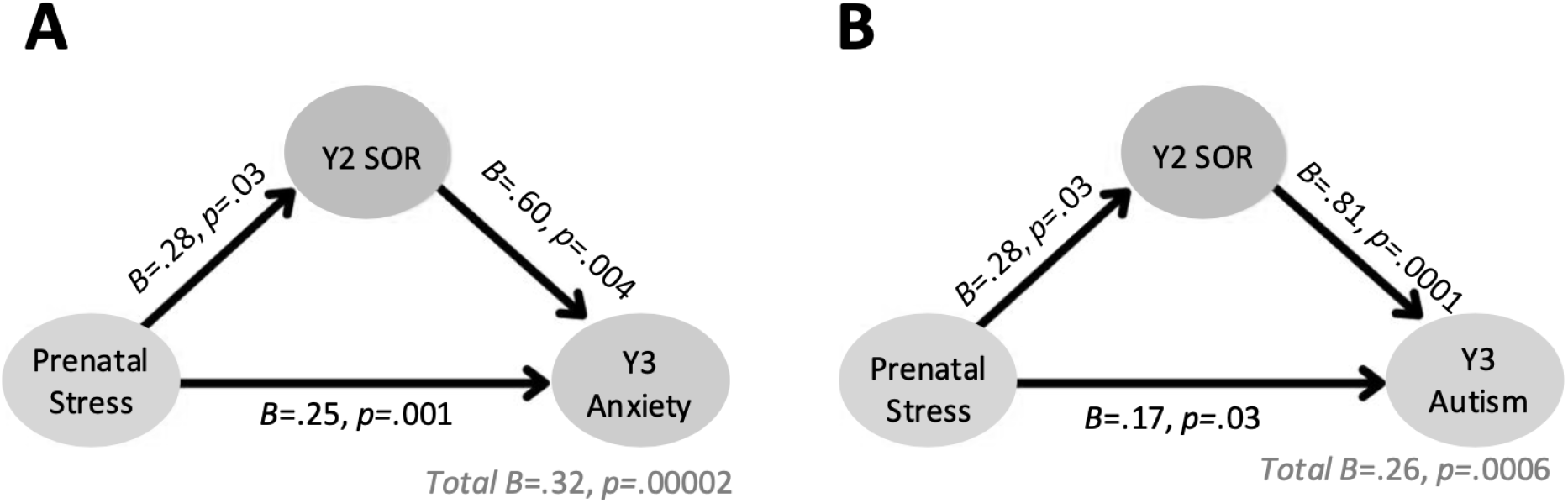
SOR at Age Two Mediates Associations of Prenatal Maternal Psychosocial Stress with Anxiety and Autism Scores at Age Three Note: (a): Path diagram of mediation model. Prenatal psychosocial stress (IV) predicts Y3 anxiety problems (DV) both directly and indirectly through Y2 SOR (mediator). Unstandardized coefficients are displayed on each path, with significance levels indicated. The indirect path through Y2 SOR was significant, indicating partial mediation by Y2 SOR. (b) Path diagram of mediation model. Prenatal psychosocial stress (IV) predicts Y3 autism-spectrum problems (DV) both directly and indirectly through Y2 SOR (mediator). Unstandardized coefficients are displayed on each path, with significance levels indicated. The indirect path through Y2 SOR was significant, indicating partial mediation by Y2 SOR.

### Functional Connectivity

Finally, to inform our understanding of the neural origins of SOR, we tested whether specific prenatal FC measures predicted infants’ emerging SOR. To maximize available sample size for this analysis, and given the strong correspondence of SOR across the Y1 and Y2 timepoints, prenatal FC was investigated in relation to a composite SOR score composed of SOR scores at the Y1 and/or Y2 timepoints. Of the nine FC pairs selected *a priori* for testing due to their associations with SOR in older children (Schwarzlose et al., 2023), infant-toddler SOR was significantly positively predicted by greater FC of the bilateral thalamus with cortical regions of 1) the somatomotor hand network, and 2) the auditory network (Figure 3; Table 4). Notably, only thalamus-somatomotor hand network FC significantly predicted infant-toddler SOR after correction for multiple comparisons. The pattern of results remained the same in posthoc sensitivity analyses and analyses that include infants born prematurely (Table S18; Table S19; Table S20). Neither early-life adversity factor was significantly related to thalamus-somatomotor hand FC, therefore we did not test whether the FC measure mediated relationships between prenatal adversity and infant-toddler SOR (see Supplement for parallel analyses in inclusive sample).

**Figure 3.**
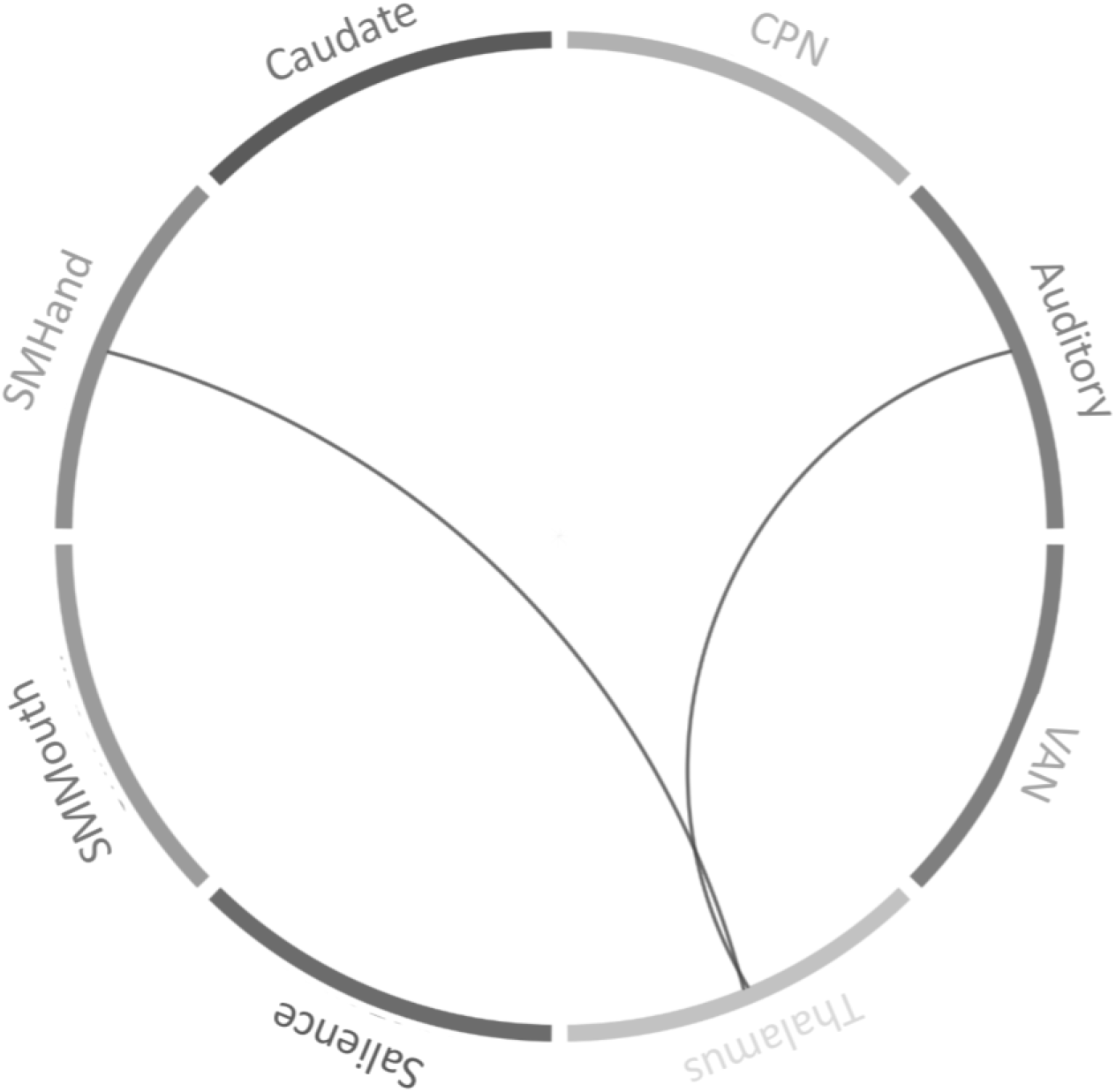
Infant-Toddler SOR is Associated with Increased Neonatal Sensory Thalamocortical FC Note: Nine a priori-selected pairs were tested altogether based on associations found in preadolescents (Schwarzlose et al, 2023). Lines denote significant, positive relationships between pairwise neonatal FC and composite SOR at one and/or two years of age before multiple comparison correction. After correction, only the relationship of SOR with FC between somatomotor hand (SMHand) parcels and the bilateral thalamus remained significant.

**Table 6.**
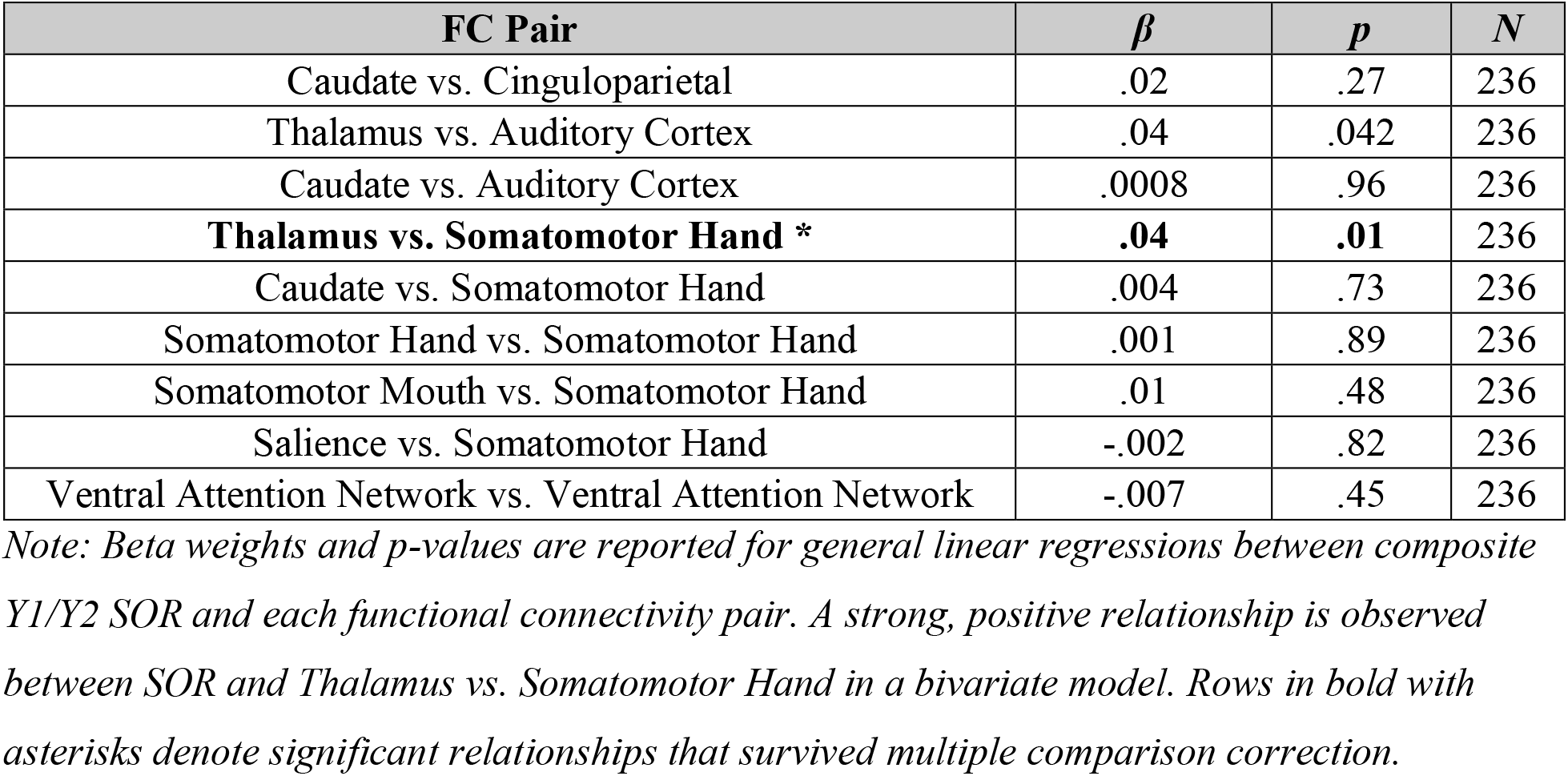
Relations between Neonatal FC and Infant-Toddler SOR.

| **FC Pair** | ***β*** | ***p*** | ***N*** |
|---|---|---|---|
| Caudate vs. Cinguloparietal | .02 | .27 | 236 |
| Thalamus vs. Auditory Cortex | .04 | .042 | 236 |
| Caudate vs. Auditory Cortex | .0008 | .96 | 236 |
| **Thalamus vs. Somatomotor Hand \*** | **.04** | **.01** | 236 |
| Caudate vs. Somatomotor Hand | .004 | .73 | 236 |
| Somatomotor Hand vs. Somatomotor Hand | .001 | .89 | 236 |
| Somatomotor Mouth vs. Somatomotor Hand | .01 | .48 | 236 |
| Salience vs. Somatomotor Hand | -.002 | .82 | 236 |
| Ventral Attention Network vs. Ventral Attention Network | -.007 | .45 | 236 |

*Note: Beta weights and p-values are reported for general linear regressions between composite Y1/Y2 SOR and each functional connectivity pair. A strong, positive relationship is observed between SOR and Thalamus vs. Somatomotor Hand in a bivariate model. Rows in bold with asterisks denote significant relationships that survived multiple comparison correction.*

## Discussion

The present work utilized a longitudinal sample enriched for early-life adversity to characterize the risk factors, neural antecedents, and clinical correlates of infant/toddler SOR, as well as its role in mediating relationships between early-life adversity and emerging neurodevelopmental and psychiatric problems. SOR at 2 years of age was positively associated with concurrent anxiety- and autism-associated behaviors, positively predicted anxiety problems and autistic traits at age 3, and predicted greater separation anxiety symptoms and diagnosis at preschool age. Although SOR at age 2 was associated with multiple forms of prenatal adversity (social disadvantage and maternal psychosocial stress), it specifically mediated relationships between prenatal psychosocial stress and emerging autism spectrum, anxiety, and internalizing problems at age 3. Infant/toddler SOR was also associated with greater neonatal functional connectivity between bilateral thalamus and sensory (i.e., somatomotor) cortical regions, implicating increased signaling in ascending sensory pathways during the first month of life in the etiology of SOR. Together these findings demonstrate the importance of infant/toddler SOR for understanding developmental pathways linking early-life adversity and sensory signaling to risk for emerging mental health challenges in early childhood.

Links between early-life adversity and psychopathology in both internalizing and externalizing domains have been found in prior literature; however, these associations have not been universally observed (Kirkbride et al., 2024) and much remains unknown about their underlying mechanisms. Previous studies using data from this unique longitudinal sample have found that prenatal SD is associated with dramatic physiological and neural effects in infancy and early childhood (Lean et al., 2022; Leverett et al., 2026; Luby et al., 2023; Nielsen et al., 2024; Triplett et al., 2022). Studies conducted with other samples have identified prenatal stress exposure as a risk factor for later child autistic traits, anxiety, and other psychiatric conditions (Beversdorf et al., 2005, 2018; Davis et al., 2011; Monk et al., 2019; Nolvi et al., 2016; Van Den Bergh et al., 2020). In the current study, we found that both prenatal exposure to SD and maternal psychosocial stress were associated with greater parent-reported SOR problems at age 2. These findings extend prior work that has shown positive associations between SOR and SD in older children (Ben-Sasson et al., 2009; Carpenter et al., 2019; Carter et al., 2011; Schwarzlose et al., 2023), providing evidence that these relationships are present in the first two years of life. Further, mediation analyses reveal that SOR at age 2 mediates associations of a specific form of ELA (prenatal maternal psychosocial stress) with emerging autism spectrum, anxiety, and internalizing problems at age 3; the same was not true for SD. Overall, this pattern of results suggests that prenatal maternal psychosocial stress may promote specific neurodevelopmental and psychiatric challenges (i.e., autistic traits and anxiety) by way of increasing children’s SOR risk. If true, this pathway could suggest new avenues for interventions at both the prenatal and infant timepoints to support adaptive sensory processing and behavioral outcomes in high-risk infants.

The study also characterized other risk factors associated with SOR in the sample. We found strong positive associations of Y1 SOR with Y2 SOR, supporting the stability across these timepoints (Baranek et al., 2019; Ben-Sasson et al., 2010; Chen et al., 2025). Notably, although premature birth is an established risk factor for SOR in prior work (e.g., Keuler et al., 2011; Van Hulle et al., 2015), we did not find that either birthweight or gestational age at birth predicted SOR at age 2 in this sample. This unexpected negative finding may reflect the fact that relatively few infants were born premature in the cohort, both because of the timing of prenatal recruitment and because the study’s intensive longitudinal design beginning in the prenatal period may have increased access to and utilization of prenatal care. We additionally note that infant sex was not a significant predictor of SOR in this sample. Other studies in child samples have found no association of SOR with sex (Ben-Sasson et al., 2009; Carson et al., 2021), association with female sex (Carpenter et al., 2019), or association with male sex (Schwarzlose et al., 2023). Therefore, additional work is needed to clarify the relationship between SOR and sex, including whether it may vary by developmental stage, SOR instrument, or informant.

Analyses of Y2 SOR in relation to concurrent and subsequent behavioral problems consistently highlighted specific associations of SOR with autism spectrum and anxiety problems, as well as to internalizing problems in general. Although clinical diagnostic measures of autism spectrum disorder were not available for this sample at time of writing, results from a clinical interview showed that SOR at age 2 was positively associated with anxiety disorder symptoms and, specifically, separation anxiety disorder diagnoses, during the preschool period. These results are consistent with prior work, primarily from studies of older children, that shows robust associations of SOR with both autistic traits and anxiety symptoms in childhood (Baranek et al., 2007; Carter et al., 2011; Conelea et al., 2014; Green et al., 2012). A recent study of data from over 15,000 school-age children supports the specificity of these associations, showing that SOR is specifically and consistently associated with autistic traits and anxiety symptoms but not symptoms of other common psychiatric conditions of childhood, when accounting for symptom co-occurrence (Luo et al., 2026). The present work demonstrated that this pattern of SOR-symptom associations is already in place within the first two years of life, suggesting that of emerging behavioral problems in association with SOR during the toddler and preschool years in this sample enriched for early-life adversity.

Finally, analysis of neuroimaging data revealed that greater neonatal thalamic FC with somatosensory cortical areas predicted greater infant/toddler SOR. This FC pair was tested because greater thalamus-somatosensory FC is associated with severe SOR in later childhood (Schwarzlose et al., 2023). This pattern of altered FC may reflect increased signaling and/or reduced gating in the ascending somatosensory signaling pathway. The present finding that these differences are in place within a month of birth, preceding the emergence of SOR and associated autism and anxiety, indicates that heightened sensory signaling in early infancy may causally contribute to these behavioral outcomes via neurodevelopmental cascades (Schwarzlose, 2025).

Strengths of the current study include the longitudinal design spanning from the prenatal to preschool timepoints and the sample enriched for early-life adversity. Limitations include the potential reporter bias due to the use of self-report measures of prenatal maternal psychosocial and parent-report measures of emerging behavioral problems, as well as reduced sample sizes at later timepoints due to participant attrition. Also, benefits from participation in the study (e.g., participant payments and prenatal medical care utilization) and exclusion of very preterm infants may affect the generalizability of some results, such as the lack of an association between gestational age at birth and later SOR. Future research in a variety of pediatric samples will be needed to fully characterize the neural bases of SOR and clarify its relation to different forms of early adversity and to emerging autistic traits and anxiety problems.

In sum, this study provides valuable insights to inform models of early adversity, sensory signaling, and psychiatric risk. These findings support the relevance of SOR to emerging behavioral problems in early life and suggest a pathway by which certain forms of early adversity, namely maternal psychosocial stress, may promote autism and anxiety. Furthermore, findings of neural differences in early infancy support models that posit a developmental cascade linking sensory signaling to SOR and mental health challenges (Schwarzlose, 2025). They highlight the pre- and perinatal period in this process, opening up new possibilities for early screening and even protective intervention by promoting the early development of adaptive sensory processing before behavioral problems emerge.

## Author Note

Stefanie Overschmidt is now at the Department of Social Work, Saint Louis University. Hexin Luo is now at the Department of Psychology, University of Wisconsin-Madison.

Deanna Barch consults for J&J. Rebecca Schwarzlose has received payments for a book first published by Mariner Books/HarperCollins and later published in translation. We thank the families that participated in the eLABE study for their time and efforts. This work was supported by the National Institute of Mental Health of the National Institutes of Health under award numbers R01MH113883 and R01MH122389, the Eunice and Kennedy Shriver National Institute of Child Health and Human Development of the National Institutes of Health under award number R00HD109454, March of Dimes, and the McDonnell Center for Systems Neuroscience. The content of this paper is solely the responsibility of the authors and does not necessarily represent the official views of the National Institutes of Health. We hereby grant to NIH, a royalty-free, nonexclusive, and irrevocable right to reproduce, publish, or otherwise use this work for Federal purposes and to authorize others to do so. This grant of rights includes the right to make the final, peer-reviewed manuscript publicly available in PubMed Central upon the Official Date of Publication.

## References

Achenbach, T. M., & Rescorla, L. A. (2000). Manual for the ASEBA preschool forms and profiles (Vol. 30). Burlington, VT: University of Vermont, Research center for children, youth ….

Alexander, B., Murray, A. L., Loh, W. Y., Matthews, L. G., Adamson, C., Beare, R., Chen, J., Kelly, C. E., Rees, S., Warfield, S. K., Anderson, P. J., Doyle, L. W., Spittle, A. J., Cheong, J. L. Y., Seal, M. L., & Thompson, D. K. (2017). A new neonatal cortical and subcortical brain atlas: The Melbourne Children’s Regional Infant Brain (M-CRIB) atlas. NeuroImage, 147, 841–851. 10.1016/j.neuroimage.2016.09.068

Baranek, G. T., Boyd, B. A., Poe, M. D., David, F. J., & Watson, L. R. (2007). Hyperresponsive Sensory Patterns in Young Children With Autism, Developmental Delay, and Typical Development. American Journal on Mental Retardation, 112(4), 233–245. 10.1352/0895-8017(2007)112%5B233:HSPIYC%5D2.0.CO;2

Baranek, G. T., Carlson, M., Sideris, J., Kirby, A. V., Watson, L. R., Williams, K. L., & Bulluck, J. (2019). Longitudinal assessment of stability of sensory features in children with autism spectrum disorder or other developmental disabilities. Autism Research, 12(1), 100–111. 10.1002/aur.2008

Ben-Sasson, A., Carter, A. S., & Briggs-Gowan, M. J. (2009). Sensory Over-Responsivity in Elementary School: Prevalence and Social-Emotional Correlates. Journal of Abnormal Child Psychology, 37(5), 705–716. 10.1007/s10802-008-9295-8

Ben-Sasson, A., Carter, A. S., & Briggs-Gowan, M. J. (2010). The Development of Sensory Over-responsivity From Infancy to Elementary School. Journal of Abnormal Child Psychology, 38(8), 1193–1202. 10.1007/s10802-010-9435-9

Ben-Sasson, A., Gal, E., Fluss, R., Katz-Zetler, N., & Cermak, S. A. (2019). Update of a Meta-analysis of Sensory Symptoms in ASD: A New Decade of Research. Journal of Autism and Developmental Disorders, 49(12), 4974–4996. 10.1007/s10803-019-04180-0

Beversdorf, D. Q., Manning, S. E., Hillier, A., Anderson, S. L., Nordgren, R. E., Walters, S. E., Nagaraja, H. N., Cooley, W. C., Gaelic, S. E., & Bauman, M. L. (2005). Timing of Prenatal Stressors and Autism. Journal of Autism and Developmental Disorders, 35(4), 471–478. 10.1007/s10803-005-5037-8

Beversdorf, D. Q., Stevens, H. E., & Jones, K. L. (2018). Prenatal Stress, Maternal Immune Dysregulation, and Their Association With Autism Spectrum Disorders. Current Psychiatry Reports, 20(9), 76. 10.1007/s11920-018-0945-4

Briggs-Gowan, M. J., & Carter, A. S. (1999). Preliminary Acceptability and Psychometrics of the Infant-Toddler Social and Emotional Assessment (ITSEA): A New Adult-Report Questionnaire. Infant Mental Health Journal: Infancy and Early Childhood, 19(4), 422–445. 10.1002/(SICI)1097-0355(199824)19:4%3C422::AID-IMHJ5%3E3.0.CO;2-U

Carpenter, K. L. H., Baranek, G. T., Copeland, W. E., Compton, S., Zucker, N., Dawson, G., & Egger, H. L. (2019). Sensory Over-Responsivity: An Early Risk Factor for Anxiety and Behavioral Challenges in Young Children. Journal of Abnormal Child Psychology, 47(6), 1075–1088. 10.1007/s10802-018-0502-y

Carson, T. B., Valente, M. J., Wilkes, B. J., & Richard, L. (2021). Brief Report: Prevalence and Severity of Auditory Sensory Over-Responsivity in Autism as Reported by Parents and Caregivers. Journal of Autism and Developmental Disorders. 10.1007/s10803-021-04991-0

Carter, A. S., Ben-Sasson, A., & Briggs-Gowan, M. J. (2011). Sensory Over-Responsivity, Psychopathology, and Family Impairment in School-Aged Children. Journal of the American Academy of Child & Adolescent Psychiatry, 50(12), 1210–1219. 10.1016/j.jaac.2011.09.010

Carter, A. S., Briggs-Gowan, M. J., Jones, S. M., & Little, T. D. (2003). The Infant-Toddler Social and Emotional Assessment (ITSEA): Factor Structure, Reliability, and Validity. Journal of Abnormal Child Psychology, 31(5), 495–514.

Cerliani, L., Mennes, M., Thomas, R. M., Di Martino, A., Thioux, M., & Keysers, C. (2015). Increased Functional Connectivity Between Subcortical and Cortical Resting-State Networks in Autism Spectrum Disorder. JAMA Psychiatry, 72(8), 767. 10.1001/jamapsychiatry.2015.0101

Chen, Y.-J., Sideris, J., Watson, L. R., Crais, E. R., & Baranek, G. T. (2022). Early developmental profiles of sensory features and links to school-age adaptive and maladaptive outcomes: A birth cohort investigation. Development and Psychopathology, 1–11. 10.1017/S0954579422001195

Chen, Y.-J., Sideris, J., Watson, L. R., Crais, E. R., & Baranek, G. T. (2025). Developmental Impacts of Early Sensory Patterns on School-Age Adaptive, Maladaptive, and Participation Outcomes in Autistic and Non-autistic Children. Journal of Autism and Developmental Disorders, 55(11), 4033–4044. 10.1007/s10803-024-06494-0

Cohen, S., Kamarck, T., & Mermelstein, R. (1994). Perceived Stress Scale. In Measuring Stress: A Guide for Health and Social Scientists (Vol. 10, pp. 1–2). Oxford University Press.

Conelea, C. A., Carter, A. C., & Freeman, J. B. (2014). Sensory Over-Responsivity in a Sample of Children Seeking Treatment for Anxiety. Journal of Developmental & Behavioral Pediatrics, 35(8), 510–521. 10.1097/DBP.0000000000000092

Cox, J. L., Holden, J. M., & Sagovsky, R. (1987). Detection of Postnatal Depression: Development of the 10-item Edinburgh Postnatal Depression Scale. British Journal of Psychiatry, 150(6), 782–786. 10.1192/bjp.150.6.782

Cummings, K. K., Lawrence, K. E., Hernandez, L. M., Wood, E. T., Bookheimer, S. Y., Dapretto, M., & Green, S. A. (2020). Sex Differences in Salience Network Connectivity and its Relationship to Sensory Over-Responsivity in Youth with Autism Spectrum Disorder. Autism Research, 13(9), 1489–1500. 10.1002/aur.2351

Davis, E. P., Glynn, L. M., Waffarn, F., & Sandman, C. A. (2011). Prenatal maternal stress programs infant stress regulation: Prenatal cortisol and infant development. Journal of Child Psychology and Psychiatry, 52(2), 119–129. 10.1111/j.1469-7610.2010.02314.x

Egger, H., Ascher, B. H., & Angold, A. (1999). Preschool Age Psychiatric Assessment (PAPA).

Gordon, E. M., Laumann, T. O., Adeyemo, B., Huckins, J. F., Kelley, W. M., & Petersen, S. E. (2016). Generation and Evaluation of a Cortical Area Parcellation from Resting-State Correlations. Cerebral Cortex, 26(1), 288–303. 10.1093/cercor/bhu239

Green, S. A., Ben-Sasson, A., Soto, T. W., & Carter, A. S. (2012). Anxiety and Sensory Over-Responsivity in Toddlers with Autism Spectrum Disorders: Bidirectional Effects Across Time. Journal of Autism and Developmental Disorders, 42(6), 1112–1119. 10.1007/s10803-011-1361-3

Helen Egger L., Angold, A., Small, B., & Copeland, W. (2020). Preschool Age Psychiatric Assessment: A Structured Parent Interview for Assessing Psychiatric Symptoms and Disorders in Preschool Children. In The Oxford Handbook of Infant, Toddler, and Preschool Mental Health Assessment, 2nd Edition (2nd ed.). Oxford University Press.

Hochberg, Y. (1988). A sharper Bonferroni procedure for multiple tests of significance. Biometrika, 75(4), 800–802.

Humphreys, K. L., & Zeanah, C. H. (2014). Deviations from the Expectable Environment in Early Childhood and Emerging Psychopathology. Neuropsychopharmacology, 40, 154–170. 10.1038/npp.2014.165

Keuler, M. M., Schmidt, N. L., Van Hulle, C. A., Lemery-Chalfant, K., & Goldsmith, H. H. (2011). Sensory Overresponsivity: Prenatal Risk Factors and Temperamental Contributions. Journal of Developmental & Behavioral Pediatrics, 32(7), 533–541. 10.1097/DBP.0b013e3182245c05

Kind, A. J. H., & Buckingham, W. R. (2018). Making Neighborhood-Disadvantage Metrics Accessible—The Neighborhood Atlas. The New England Journal of Medicine, 378(26). 10.1056/NEJMp1802313

Kirkbride, J. B., Anglin, D. M., Colman, I., Dykxhoorn, J., Jones, P. B., Patalay, P., Pitman, A., Soneson, E., Steare, T., Wright, T., & Griffiths, S. L. (2024). The social determinants of mental health and disorder: Evidence, prevention and recommendations. World Psychiatry, 23(1), 58–90. 10.1002/wps.21160

Lean, R. E., Smyser, C. D., Brady, R. G., Triplett, R. L., Kaplan, S., Kenley, J. K., Shimony, J. S., Smyser, T. A., Miller, J. P., Barch, D. M., Luby, J. L., Warner, B. B., & Rogers, C. E. (2022). Prenatal exposure to maternal social disadvantage and psychosocial stress and neonatal white matter connectivity at birth. Proceedings of the National Academy of Sciences, 119(42), e2204135119. 10.1073/pnas.2204135119

Leverett, S. D., Poolos, O., Brady, R. G., Tillman, R., Lean, R. E., Gerstein, E. D., Anaya, B., Triplett, R. L., Alexopoulos, D., Warner, B., Luby, J. L., Smyser, C. D., Rogers, C. E., & Barch, D. M. (2026). Neonatal Brain Volumes and Early Parenting Behavior as Mediators in Associations Between Prenatal Social Disadvantage and Socioemotional Problems in Toddlers. Biological Psychiatry, 99(5), 387–397. 10.1016/j.biopsych.2025.05.015

Levy, S. E., Rescorla, L. A., Chittams, J. L., Kral, T. J., Moody, E. J., Pandey, J., Pinto-Martin, J. A., Pomykacz, A. T., Ramirez, A., Reyes, N., Rosenberg, C. R., Schieve, L., Thompson, A., Young, L., Zhang, J., & Wiggins, L. (2019). ASD Screening with the Child Behavior Checklist/1.5-5 in the Study to Explore Early Development. Journal of Autism and Developmental Disorders, 49(6), 2348–2357. 10.1007/s10803-019-03895-4

Lewis, T. T., Yang, F. M., Jacobs, E. A., & Fitchett, G. (2012). Racial/Ethnic Differences in Responses to the Everyday Discrimination Scale: A Differential Item Functioning Analysis. American Journal of Epidemiology, 175(5), 391–401.

Luby, J. L., England, S. K., Barch, D. M., Warner, B. B., Rogers, C., Smyser, C. D., Triplett, R., Arora, J., Smyser, T. A., Slavich, G. M., Zhao, P., Stout, M., Herzog, E., & Miller, J. P. (2023). Social disadvantage during pregnancy: Effects on gestational age and birthweight. Journal of Perinatology, 43(4), 477–483. 10.1038/s41372-023-01643-2

Luo, H., Kim, A. W., Abbacchi, A. M., Constantino, J. N., Gurnett, C. A., Luby, J. L., Perino, M. T., Barch, D. M., Sylvester, C. M., Camacho, M. C., & Schwarzlose, R. F. (2026). Specific, Replicable Behavioral and Neural Correlates of Sensory Over-Responsivity in Childhood. Journal of the American Academy of Child & Adolescent Psychiatry, S0890856726015698. 10.1016/j.jaac.2026.08.003

McCormick, C., Hepburn, S., Young, G. S., & Rogers, S. J. (2016). Sensory symptoms in children with autism spectrum disorder, other developmental disorders and typical development: A longitudinal study. Autism, 20(5), 572–579. 10.1177/1362361315599755

McLaughlin, K. A., Green, J. G., Gruber, M. J., Sampson, N. A., Zaslavsky, A. M., & Kessler, R. C. (2012). Childhood Adversities and First Onset of Psychiatric Disorders in a National Sample of US Adolescents. JAMA Psychiatry, 69(11), 1151–1160. doi:10.1001/archgenpsychiatry.2011.2277

Menyhart, O., Weltz, B., & Győrffy, B. (2021). MultipleTesting.com: A tool for life science researchers for multiple hypothesis testing correction. PLOS ONE, 16(6), e0245824. 10.1371/journal.pone.0245824

Monk, C., Lugo-Candelas, C., & Trumpff, C. (2019). Prenatal Developmental Origins of Future Psychopathology: Mechanisms and Pathways. Annual Review of Clinical Psychology, 15(1), 317–344. 10.1146/annurev-clinpsy-050718-095539

Nair, A., Jalal, R., Liu, J., Tsang, T., McDonald, N. M., Jackson, L., Ponting, C., Jeste, S. S., Bookheimer, S. Y., & Dapretto, M. (2021). Altered Thalamocortical Connectivity in 6-Week-Old Infants at High Familial Risk for Autism Spectrum Disorder. Cerebral Cortex, 31(9), 4191–4205. 10.1093/cercor/bhab078

Narvekar, N., Carter Leno, V., Pasco, G., Begum Ali, J., Johnson, M. H., Charman, T., Jones, E. J. H., & The STAARS Team. (2024). The roles of sensory hyperreactivity and hyporeactivity in understanding infant fearfulness and emerging autistic traits. Journal of Child Psychology and Psychiatry, jcpp.13941. 10.1111/jcpp.13941

National Institute of Health Epidemiology and Genomics Research Program. (2010). Diet History Questionnaire, version 2.0 (DHQ-II). SNAP Education Connection.

Nelson, C. A., Sullivan, E. F., & Valdes, V. (2025). Early Adversity Alters Brain Architecture and Increases Susceptibility to Mental Health Disorders. Nature Reviews Neuroscience, 26, 642–656. 10.1038/s41583-025-00948-9

Nielsen, A. N., Triplett, R. L., Bernardez, L. M., Tooley, U. A., Herzberg, M. P., Lean, R. E., Kaplan, S., Meyer, D., Kenley, J. K., Alexopoulos, D., Losielle, D., Latham, A., Smyser, T. A., Agrawal, A., Shimony, J. S., Jackson, J. J., Miller, J. P., Raichle, M. E., Warner, B. B., … Smyser, C. D. (2024). Prenatal social disadvantage is associated with alterations in functional networks at birth. Proceedings of the National Academy of Sciences, 121(50), e2405448121. 10.1073/pnas.2405448121

Nolvi, S., Karlsson, L., Bridgett, D. J., Korja, R., Huizink, A. C., Kataja, E.-L., & Karlsson, H. (2016). Maternal prenatal stress and infant emotional reactivity six months postpartum. Journal of Affective Disorders, 199, 163–170. 10.1016/j.jad.2016.04.020

Peverill, M., Dirks, M. A., Narvaja, T., Herts, K. L., Comer, J. S., & McLaughlin, K. A. (2021). Socioeconomic status and child psychopathology in the United States: A meta-analysis of population-based studies. Clinical Psychology Review, 83. 10.1016/j.cpr.2020.101933

Reiss, F. (2013). Socioeconomic Inequalities and Mental Health Problems in Children and Adolescents: A Systematic Review. Social Science & Medicine, 90, 24–31. 10.1016/j.socscimed.2013.04.026

Rescorla, L. A., Winder-Patel, B. M., Paterson, S. J., Pandey, J., Wolff, J. J., Schultz, R. T., & Piven, J. (2019). Autism spectrum disorder screening with the CBCL/1½–5: Findings for young children at high risk for autism spectrum disorder. Autism, 23(1), 29–38. 10.1177/1362361317718482

Schwarzlose, R. F. (2025). Infant Sensory Gating and a Developmental Cascade to Autistic Traits and Anxiety. Neuropsychopharmacology. 10.1038/s41386-025-02253-6

Schwarzlose, R. F., Tillman, R., Hoyniak, C. P., Luby, J. L., & Barch, D. M. (2023). Sensory Over-responsivity: A Feature of Childhood Psychiatric Illness Associated With Altered Functional Connectivity of Sensory Networks. Biological Psychiatry, 93(1), 92–101. 10.1016/j.biopsych.2022.09.004

Slavich, G. M., & Shields, G. S. (2018). Assessing Lifetime Stress Exposure Using the Stress and Adversity Inventory for Adults (Adult STRAIN): An Overview and Initial Validation. Psychosomatic Medicine, 80(1), 17–27. 10.1097/PSY.0000000000000534

Tooley, U. A., Kenley, J. K., Camacho, M. C., Latham, A., Alexopoulos, D., Nielsen, A. N., Smyser, T. A., Warner, B. B., Shimony, J. S., Neil, J. J., Luby, J. L., Barch, D. M., Rogers, C. E., & Smyser, C. D. (2025). Structure–function coupling in the first month of life: Associations with age and attention. Proceedings of the National Academy of Sciences, 122(23), e2412729122. 10.1073/pnas.2412729122

Triplett, R. L., Lean, R. E., Parikh, A., Miller, J. P., Alexopoulos, D., Kaplan, S., Meyer, D., Adamson, C., Smyser, T. A., Rogers, C. E., Barch, D. M., Warner, B., Luby, J. L., & Smyser, C. D. (2022). Association of Prenatal Exposure to Early-Life Adversity With Neonatal Brain Volumes at Birth. JAMA Network Open, 5(4), e227045. 10.1001/jamanetworkopen.2022.7045

Tsang, T., Green, S. A., Liu, J., Lawrence, K., Jeste, S., Bookheimer, S. Y., & Dapretto, M. (2024). Salience network connectivity is altered in 6-week-old infants at heightened likelihood for developing autism. Communications Biology, 7(1), 485. 10.1038/s42003-024-06016-9

Tseliou, F., Collishaw, S., Price, A., & Sumner, P. (2025). Sensory hyperacusis as a predictor of anxiety in adolescence. Journal of Child Psychology and Psychiatry. 10.1111/jcpp.70027

Tu, J. C., Wang, Y., Wang, X., Dierker, D., Sobolewski, C. M., Day, T. K. M., Kardan, O., Miranda-Domínguez, Ó., Moore, L. A., Feczko, E., Fair, D. A., Elison, J. T., Gordon, E. M., Laumann, T. O., Eggebrecht, A. T., & Wheelock, M. D. (2025). A subset of cortical areas exhibit adult-like functional network patterns in early childhood. Developmental Cognitive Neuroscience, 73, 101551. 10.1016/j.dcn.2025.101551

Vaidya, N., Marquand, A. F., Nees, F., Siehl, S., & Schumann, G. (2024). The Impact of Psychosocial Adversity on Brain and Behaviour: An Overview of Existing Knowledge and Directions for Future Research. Molecular Psychiatry, 29, 3245–3267. 10.1038/s41380-024-02556-y

Van Den Bergh, B. R. H., Van Den Heuvel, M. I., Lahti, M., Braeken, M., De Rooij, S. R., Entringer, S., Hoyer, D., Roseboom, T., Räikkönen, K., King, S., & Schwab, M. (2020). Prenatal developmental origins of behavior and mental health: The influence of maternal stress in pregnancy. Neuroscience & Biobehavioral Reviews, 117, 26–64. 10.1016/j.neubiorev.2017.07.003

Van Hulle, C., Lemery-Chalfant, K., & Goldsmith, H. H. (2015). Trajectories of Sensory Over-Responsivity from Early to Middle Childhood: Birth and Temperament Risk Factors. PLOS ONE, 10(6), e0129968. 10.1371/journal.pone.0129968

Venables, W., & Ripley, B. (2002). Modern Applied Statistics with S (4th ed.). Springer. https://www.stats.ox.ac.uk/pub/MASS4/

Wagner, L., Banchik, M., Okada, N. J., McDonald, N., Jeste, S. S., Bookheimer, S. Y., Green, S. A., & Dapretto, M. (2023). Associations between thalamocortical functional connectivity and sensory over-responsivity in infants at high likelihood for ASD. Cerebral Cortex, 33(12), 8075–8086. 10.1093/cercor/bhad100

Wickham, H., Francois, R., Henry, L., Muller, K., & Vaughan, D. (2025). dplyr: A Grammar of Data Manipulation (Version 1.1.4) [Computer software]. https://dplyr.tidyverse.org

