## Supplemental Materials for "Infant Sensory Over-Responsivity: Relations to Early Life Adversity, Emerging Behavioral Problems, and Sensory Functional Connectivity"

**Supplemental Methods**

<u>Adversity Measures</u>

The SD latent factor is derived from maternal health insurance status, highest maternal level of educational attainment, household Income-to-Needs Ratio in each trimester, national Area Deprivation Index percentiles (Kind & Buckingham, 2018), and Healthy Eating Index(Guenther et al., 2014). The psychosocial stress (PSS) latent factor is derived from maternal scores from the Edinburgh Postnatal Depression Scale in each trimester (Cox et al., 1987), Cohen Perceived Stress Scale in each trimester (Cohen et al., 1983), Stress and Adversity Inventory for Adults lifetime count and severity at the neonatal scan or follow-up at one or two years (Slavich & Shields, 2018), and the Everyday Discrimination Scale at the neonatal timepoint (Lewis et al., 2012).

<u>Child Behavioral Measures</u>

The ITSEA is a parent-report measure assessing social-emotional functioning in 12- to 36-month-old children (Carter et al., 1999, 2003). The Sensory Sensitivity subscale is computed as the average raw score across seven items assessing children's negative responses to common sensory stimuli, including touch, sound, smell, sight, and motion. Responses are collected with a three-point Likert scale: *Rarely/Not True of My Child (0); Sometimes/Somewhat True of My Child (1); and Often/Very True of My Child (2).*

The CBCL 1.5-5 is a 99-item parent-report measure of emotional and behavioral problems in toddlers and preschool-age children. Items are scored on a three-point Likert scale: *Not true (as far as you know) (0); Somewhat or sometimes true (1); and Very true or often true (2).* All scores are computed as the sum of individual item responses (Achenbach & Rescorla, 2000).

The second-edition Preschool Age Psychiatric Assessment (PAPA) is designed to assess DSM-V psychiatric symptoms and their effect on a child's social functioning in preschool-aged children between two and eight years of age (Helen Egger L. et al., 2020). Diagnoses are determined based on specific criteria from the developers of the interview (Egger et al., 1999).

<u>Negative Binomial Model</u>

Negative binomial models were used in behavioral analyses to account for a zero-skewed distribution in behavioral outcome variables. Although zero-inflated models are also common when analyzing zero-skewed data, we opted for the negative binomial due to the model accounting more accurately for data in which zero is part of a continuum of scores, whereas a zero-inflated model is more fitting for use with data in which a zero value is categorically different than all other non-zero scores.

<u>Composite SOR Measure</u>

To create the composite Y1/Y2 SOR measure, both paired and independent t-tests were performed to evaluate the difference in means between Y1 SOR and Y2 SOR scores. Both paired and independent t-tests revealed that Y1 SOR scores were lower on average than Y2 SOR scores ($t = -2.33$, $p = 0.02$; $t = -2.26$, $p = 0.02$). The difference between the average Y2 SOR score and average Y1 SOR score was then computed, and this difference was added to the Y1 SOR scores to create an adjusted measure. Either adjusted Y1 SOR or Y2 SOR data was used if a child had a measurement at only one timepoint, whereas adjusted Y1 SOR and Y2 SOR were averaged to create a single score for children with data at both timepoints.

**Supplemental Results**

Gestational Age and SOR at 2 Years of Age

Despite prior evidence showing a negative relationship between gestational age (GA) at birth and later SOR, our analyses did not yield similar results; in fact, we found no relationship between the two measures. Follow-up analyses using a full unwinsorized sample sought to establish a step-function between gestational age and Y2 SOR (see Figure S1b); however, no evidence was found to indicate this relationship. The inquiry did, however, yield evidence that both gestational age and Y2 SOR measures have a finite number of values, causing multiple data points to overlap. Furthermore, more infants were born at a greater gestational age, such that a wider range of SOR scores could be observed at the higher end of the GA spectrum (see Figure S1a).

Additional analyses sought to understand how the central tendency of the relationship between GA and Y2 SOR changed across the spectrum of GA in the sample. Since our sample is enriched for social disadvantage, the relationship between GA and Y2 SOR was examined across social disadvantage groups by dichotomizing the prenatal social disadvantage latent factor into high and low SD groups; however, we found a lack of directionality in these relationships. Although a slight, positive relationship between GA and Y2 SOR was observed in the low social disadvantage group, the observed results were not notably different than those found in prior analyses, leading to the conclusion that the sample did not contain enough premature babies to reliably estimate SOR's relationship to GA in the current work.

Predictors of SOR at 2 Years of Age

Supplemental models of prenatal social disadvantage, maternal psychosocial stress, and Y1 SOR predicting Y2 SOR each included age and, separately, sex as covariates. Prenatal social disadvantage remained significantly associated with SOR at two years of age when accounting for child sex but not age at year two (see Table S5). Maternal psychosocial stress remained significantly associated with SOR at two years of age when accounting for year two age and child sex (see Table S5). Continuity of SOR across timepoints was also conserved with addition of age at Y2 and sex (see Table S5). Birthweight, gestational age at birth, and child sex were not tested with inclusion of additional covariates, as they were not found to be significant in initial analyses.

Parallel models were performed using only infants at or above 30 weeks gestational age for analyses. SD and maternal psychosocial stress were each significantly associated with Y2 SOR (see Table S6). By contrast, birthweight, gestational age at birth, and child sex were not significant predictors of Y2 SOR (see Table S6). Analyses indicate continuity of SOR across infant timepoints, such that Y1 SOR significantly predicted Y2 SOR (see Table S6). Follow-up analyses confirmed that results remained the same after inclusion of key covariates (i.e., child sex and age; see Table S6).

Relations between SOR and Emerging Symptoms

Supplemental models of Y2 SOR predicting Y2 CBCL internalizing and externalizing problem scales, as well as DSM-oriented problem scales (i.e., affective problems, anxiety problems, autism spectrum problems, ADHD problems), each separately included prenatal social disadvantage, age at year two, and sex. Only relationships found to be significant in bivariate analyses were modeled with these additional covariates. Both concurrent internalizing problems and externalizing problems remained significant after inclusion of prenatal social disadvantage, age at year two, and child sex (see Table S7). Affective problems, anxiety problems, autism spectrum problems, and ADHD problems remained significant after inclusion of prenatal social disadvantage, age at year two, and child sex (see Table S7).

Parallel models were performed using a more inclusive sample of infants born at or above 30 weeks gestational age for analyses. Y2 SOR was significantly associated with both concurrent internalizing problems and externalizing problems (see Table S8). Associations remained significant in models that separately included key covariates (i.e., SD, age, sex; see Tables S8 and S9). Among DSM-oriented subscale scores, SOR was related to greater concurrent anxiety problems, autism spectrum problems, and affective problems (see Table S8). Additional analyses confirmed that the observed effects remained significant after inclusion of key covariates (i.e., SD, age, sex; see Tables S8 and S9).

Additional models probing the relationship between Y2 SOR and Y3 CBCL internalizing and externalizing problem scales and DSM-oriented scales (i.e., affective problems, anxiety problems, autism spectrum problems, and ADHD problems) each separately included prenatal social disadvantage, age at year two, age at year three, and sex. Only relationships found to be significant in bivariate analyses were modeled with these additional covariates. Internalizing problems and externalizing problems remained significant after inclusion of prenatal social disadvantage, age at year two, age at year three, and child sex (see Tables S10 and S11). Analyses of SOR and key covariates with DSM-oriented subscale scores revealed significant associations with affective problems, anxiety problems, and autism spectrum problems after inclusion of prenatal social disadvantage, age at year two, age at year three, and child sex, while ADHD problems survived inclusion of only age at year two, age at year three, and child sex (see Tables S10 and Table S11).

Parallel models were performed using a sample of infants born at or above 30 weeks gestational age for analyses. Y2 SOR significantly predicted Y3 internalizing problems, anxiety problems, and autism spectrum problems (see Table S12). All effects survived inclusion of Y2 problem scores, SD, age at year two, age at year three, and sex as a covariates in independent models (see Tables S12 and S13).

SOR at age 2 predicts preschool symptoms in clinical interview

Parallel models were performed using a sample of infants born at or above 30 weeks gestational age for analyses. In analyses of Y2 SOR predicting number of psychiatric symptoms assessed at follow-up timepoints when children were between four and six years of age, Y2 SOR was significantly associated with greater number of separation anxiety symptoms but not internalizing symptoms, externalizing symptoms, social phobia symptoms, or generalized anxiety symptoms; this effect survived inclusion of age at PAPA administration as a covariate (see Table S15). Analyses of Y2 SOR predicting categorical preschool diagnoses found no significant results (see Table S17).

Mediation Analyses

Given the significant relationships found between prenatal maternal psychosocial stress and both Y3 internalizing and externalizing problems, we sought to explore the mediating role of Y2 SOR in these relationships. In mediation analyses on data from 164 children, Y2 SOR partially mediated the relationship between PSS and Y3 internalizing problems (see Figure S5a). The direct effect of PSS on Y3 anxiety remained significant. Y2 SOR did not mediate the relationship of PSS and Y3 externalizing problems, despite the direct effect of PSS on Y3 externalizing problems remaining significant (see Figure S5b).

In parallel mediation analyses on a sample of infants born at or above 30 weeks gestational age ($n = 186$), paths from PSS to anxiety and autism spectrum problems, as well as from SD to anxiety problems, were explored. Y2 SOR partially mediated the relationship between PSS and Y3 anxiety (see Figure S6a). The direct effect of PSS on Y3 anxiety remained significant. Y2 SOR also partially mediated the relationship between PSS and Y3 autism spectrum problems (see Figure S6b). The direct effect of PSS on Y3 autism spectrum problems remained significant. Additionally, the relationship between SD and Y3 anxiety was fully mediated by Y2 SOR (see Figure S6c). The direct effect of SD on Y3 anxiety became non-significant. A mediation including SD and Y3 autism spectrum problems was not conducted due to a lack of significant relation between the measures ($\beta = 0.07$, $p = 0.30$).

Further mediation analyses were performed for paths from PSS to both Y3 internalizing and externalizing problems. Y2 SOR partially mediated the relationship between PSS and Y3 internalizing problems (see Figure S7a). The direct effect of PSS on Y3 internalizing problems remained significant. The relationship between PSS and Y3 externalizing problems was not, however, mediated by Y2 SOR, despite the direct effect of PSS on Y3 externalizing problems remaining significant (see Figure S7b). Mediation analyses including SD and Y3 internalizing and externalizing problems were not conducted due to a lack of significant relation between the measures ($\beta = 0.12$, $p = 0.052$; $\beta = 0.065$, $p = 0.23$).

Functional Connectivity Analyses

Supplemental models of a composite score of Y1 and Y2 SOR predicting neonatal network-level functional connectivity were conducted in models found to be significant in bivariate analyses. These models separately included number of retained frames, PSS, and SD, with thalamus-somatomotor hand FC remaining significant in all post-hoc models. Additional analyses of thalamus-auditory cortex FC only yielded significant results when including number of frames retained but not SD or PSS (see Table S18).

Parallel models were performed using a larger sample of infants born at or above 30 weeks gestational age. Infant-toddler SOR was significantly associated with greater FC between thalamus and cortical regions of the somatomotor hand network and greater FC between thalamus and auditory cortex (see Table S19). These significant associations were also tested in multivariate analyses separately including frames retained, social disadvantage, and maternal psychosocial stress. The association between SOR and thalamus-auditory cortex only remained significant after including number of frames retained but not after inclusion of SD or PSS, while the association between SOR and thalamus-somatomotor hand network remained significant only after inclusion of number of frames retained and PSS but not SD (see Table S20).

A parallel mediation analysis to test the mediating effect of Y0 thalamus-somatomotor hand FC on the relationship between prenatal SD and Y1/Y2 composite SOR was performed ($n = 275$). SD significantly predicted both Y1/Y2 SOR and Y0 thalamus-somatomotor hand FC in separate bivariate models ($\beta = 0.28$, $p < 0.01$; $\beta = 0.008$, $p = 0.02$); however, in a multivariate model of SD and thalamus-somatomotor hand FC predicting Y1/Y2 SOR, thalamus-somatomotor hand FC did not significantly predict Y1/Y2 SOR ($\beta = 1.19$, $p = 0.49$). The direct effect of SD on Y1/Y2 SOR did remain significant ($\beta = 0.27$, $p = 0.01$). The mediating effect of thalamus-somatomotor hand FC on the relationship between PSS and Y1/Y2 SOR was not tested due to a lack of significant association between PSS and thalamus-somatomotor hand FC ($\beta = 0.004$, $p = 0.25$).

**Supplemental Tables**

**Table S1. Number of valid datapoints.** *Number of valid datapoints collected for each measure in the sample (adjusted to include only infants born at or above 37 weeks gestational age).*

| Measure | Number of Datapoints |
|---|---|
| Prenatal Social Disadvantage | 342 |
| Prenatal Psychosocial Stress | 342 |
| Birthweight | 343 |
| Gestational Age at Birth | 343 |
| Child Sex | 343 |
| Year One ITSEA | 256 |
| Year Two ITSEA | 235 |
| Year Two CBCL | 142 |

| | |
|---|---|
| Year Three CBCL | 189 |
| Preschool-Age PAPA | 221 |
| Neonatal Functional Connectivity | 319 |

**Table S2. ITSEA Sensory Sensitivity Subscale.** *The below items comprise the ITSEA sensory sensitivity subscale used for Y1 and Y2 SOR variables.*

| ***These items were scored on a three-point Likert scale (0 = Rarely/Never, 2 = Often):*** |
|---|
| Won't touch some objects because of how they feel |
| Bothered by loud noises or bright lights |
| Dislikes some foods because of how they feel |
| Bothered by certain odors (smells) |
| Bothered by how some things feel on his or her skin (for example, clothing seams, certain fabrics, etc.) |
| Easily startled |
| Bothered by being in motion (for example, swinging, spinning, being tossed in the air, or bouncing) |

**Table S3. CBCL 1.5-5 DSM-Oriented Scales.** *The below items comprise the five DSM-oriented subscales used for Y2 and Y3 symptom variables.*

| ***These items were scored on a three-point Likert scale (0 = Not True as far as you know, 2 = Very True or Often True):*** |
|---|
| Affective Problems: |
| Cries a lot |
| Doesn't eat well |
| Has trouble getting to sleep |
| Looks unhappy without good reason |
| Overeating |
| Overtired |
| Shows little interest in things around them |
| Sleeps less than most kids during day and/or night |
| Underactive, slow moving, or lacks energy |
| Unhappy, sad or depressed |
| Anxiety Problems: |
| Clings to adults or too dependent |
| Doesn't want to sleep alone |
| Doesn't want to go out of home |
| Fears certain animals, situations, or places |
| Gets too upset when separated from parents |
| Nervous, highstrung, or tense |
| Nightmares |
| Shows panic for no good reason |
| Too fearful or anxious |

| |
|---|
| Worries |
| Pervasive Developmental Problems: |
| Afraid to try new things |
| Avoids looking others in the eye |
| Can’t stand having things out of place |
| Disturbed by any change in routine |
| Doesn’t answer when people talk to them |
| Doesn’t get along with other children |
| Repeatedly rocks head or body |
| Seems unresponsive to affection |
| Shows little affection toward people |
| Speech problem |
| Strange behavior |
| Upset by new people or situations |
| Withdrawn, doesn’t get involved with others |
| Attention Deficit/Hyperactivity Problems: |
| Can’t concentrate, can’t pay attention for long |
| Can’t sit still, restless, or hyperactive |
| Can’t stand waiting, wants everything now |
| Demands must be met immediately |
| Gets into everything |
| Quickly shifts from one activity to another |
| Oppositional Defiant Problems: |
| Defiant |
| Disobedient |
| Angry moods |
| Stubborn, sullen, or irritable |
| Temper tantrums or hot temper |
| Uncooperative |

**Table S4. Regression analysis summary.** *Full list of all models run both in main cohort of children born at or above 37 weeks gestational age. Outcome variables in parentheses were analyzed only in mediation analyses. Similar analyses were run in a more inclusive cohort of children born at or above 30 weeks gestational age. One asterisk denotes models run only in the cohort of children born at or above 37 weeks gestational age, and two asterisks denotes models run only in the main cohort of children born at or above 30 weeks gestational age.*

| Predictor Variable | Number of Models | Outcome Variables |
|---|---|---|
| SD | 5 | Y2 SOR (Y3 Anxiety, Y3 PDD, Y3 Internalizing, Y3 Externalizing, Composite SOR, Y0 Thalamus vs. SMH) |
| *Y2 Age Covariate* | | |
| *SD* | *1* | *Y2 SOR* |

| | | |
|---|---|---|
| *Sex Covariate* | | |
| *SD* | *1* | *Y2 SOR* |
| *Y2 SOR Covariate* | | |
| *SD* | *1* | *(Y3 Anxiety)*** |
| *Y0 Thalamus vs. SMH Covariate* | | |
| *SD* | *1* | *(Composite SOR)*** |
| PSS | | Y2 SOR (Y3 Anxiety, Y3 PDD, Y3 Internalizing, Y3 Externalizing, Composite SOR, Y0 Thalamus vs. SMH) |
| *Y2 Age Covariate* | | |
| *PSS* | *1* | *Y2 SOR* |
| *Sex Covariate* | | |
| *PSS* | *1* | *Y2 SOR* |
| *Y2 SOR Covariate* | | |
| *PSS* | *4* | *(Y3 Anxiety, Y3 PDD, Y3 Internalizing, Y3 Externalizing)* |
| Birthweight | 1 | Y2 SOR |
| Gestational Age | 1 | Y2 SOR |
| Sex | 1 | Y2 SOR |
| Y1 SOR | 1 | Y2 SOR |
| *Y2 Age Covariate* | | |
| *Y1 SOR* | *1* | *Y2 SOR* |
| *Sex Covariate* | | |
| *Y1 SOR* | *1* | *Y2 SOR* |
| Y2 SOR | 24 | Y2 Internalizing, Y2 Externalizing, Y2 Anxiety, Y2 Affective, Y2 PDD, Y2 ADHD, Y2 ODD, Y3 Internalizing, Y3 Externalizing, Y3 Affective, Y3 Anxiety, Y3 PDD, Y3 ADHD, Y3 ODD, PAPA Number of Internalizing Symptoms, PAPA Number of Externalizing Symptoms, PAPA Number of Separation Anxiety Symptoms, PAPA Number of Social Phobia Symptoms, PAPA Number of GAD Symptoms, PAPA Any Internalizing Disorder Diagnosis, PAPA Any Externalizing Disorder Diagnosis, PAPA Separation Anxiety Diagnosis, PAPA Social Phobia Diagnosis, PAPA GAD Diagnosis |
| *SD Covariate* | | |
| *Y2 SOR* | *12* | *Y2 Internalizing, Y2 Externalizing, Y2 Anxiety, Y2 Affective, Y2 PDD, Y2 ADHD*, Y3 Internalizing, Y3 Externalizing*, Y3 Affective*, Y3 Anxiety, Y3 PDD, Y3 ADHD** |
| *Y2 Age Covariate* | | |

| | | |
|---|---|---|
| *Y2 SOR* | *12* | *Y2 Internalizing, Y2 Externalizing, Y2 Anxiety, Y2 Affective, Y2 PDD, Y2 ADHD*, Y3 Internalizing, Y3 Externalizing*, Y3 Affective*, Y3 Anxiety, Y3 PDD, Y3 ADHD** |
| *Sex Covariate* | | |
| *Y2 SOR* | *12* | *Y2 Internalizing, Y2 Externalizing, Y2 Anxiety, Y2 Affective, Y2 PDD, Y2 ADHD*, Y3 Internalizing, Y3 Externalizing*, Y3 Affective*, Y3 Anxiety, Y3 PDD, Y3 ADHD** |
| *Y3 Age Covariate* | | |
| *Y2 SOR* | *6* | *Y3 Internalizing, Y3 Externalizing*, Y3 Affective*, Y3 Anxiety, Y3 PDD, Y3 ADHD** |
| *Y2 Internalizing Covariate* | | |
| *Y2 SOR* | *1* | *Y3 Internalizing* |
| *Y2 Externalizing Covariate* | | |
| *Y2 SOR* | *1* | *Y3 Externalizing** |
| *Y2 Affective Covariate* | | |
| *Y2 SOR* | *1* | *Y3 Affective** |
| *Y2 Anxiety Covariate* | | |
| *Y2 SOR* | *1* | *Y3 Anxiety* |
| *Y2 PDD Covariate* | | |
| *Y2 SOR* | *1* | *Y3 PDD* |
| *Y2 ADHD Covariate* | | |
| *Y2 SOR* | *1* | *Y3 ADHD** |
| *PAPA Age* | | |
| *Y2 SOR* | *6* | *PAPA Number of Internalizing Symptoms*, PAPA Number of Separation Anxiety Symptoms, PAPA Number of Social Phobia Symptoms*, PAPA Number of GAD Symptoms*, PAPA Separation Anxiety Diagnosis*, PAPA Social Phobia Diagnosis** |
| Composite SOR | 9 | Y0 Caudate vs. CP, Y0 Thalamus vs. Auditory, Y0 Caudate vs. Auditory, Y0 Thalamus vs. SMH, Y0 Caudate vs. SMH, Y0 SMH vs. SMH, Y0 SMM vs. SMH, Y0 Salience vs. SMH, Y0 VAN vs. VAN |
| *SD Covariate* | | |
| *Composite SOR* | *2* | *Y0 Thalamus vs. Auditory, Y0 Thalamus vs. SMH* |
| *PSS Covariate* | | |
| *Composite SOR* | *2* | *Y0 Thalamus vs. Auditory, Y0 Thalamus vs. SMH* |
| *Number of Frames Retained Covariate* | | |

| | | |
|---|---|---|
| *Composite SOR* | *2* | *Y0 Thalamus vs. Auditory, Y0 Thalamus vs. SMH* |

**Table S5. Early life adversity predicts SOR at age two.** *Early life adversity and demographic variables in relation to SOR at two years of age after inclusion of key covariates (age at Y2 and sex included in independent models). The table reports unstandardized beta weights and p-values for negative binomial regression models of Y2 SOR by individual risk variable. Statistically significant effects are shown in bold.*

| | **Y2 age included as covariate** | | | **Sex included as covariate** | | |
|---|---|---|---|---|---|---|
| **Predictor** | **Beta Weight** | **P-Value** | **N** | **Beta Weight** | **P-Value** | **N** |
| **Prenatal Social Disadvantage** | 0.30 | 0.07 | 142 | **0.30** | **0.01** | 235 |
| **Prenatal Maternal Psychosocial Stress** | **0.34** | **0.02** | 142 | **0.30** | **0.005** | 235 |
| Birthweight | - | - | - | - | - | - |
| Gestational Age at Birth | - | - | - | - | - | - |
| Sex | - | - | - | - | - | - |
| **Sensory Over-Responsivity at One Year of Age** | **1.39** | **0.02** | 113 | **1.42** | **0.0007** | 197 |

**Table S6. Early life adversity predicts SOR at age two in inclusive sample.** *Early life adversity and demographic variables in relation to SOR at two years of age using sample of infants born at or after 30 weeks gestational age and after inclusion of key covariates (age at Y2 and sex included in independent models). The table reports unstandardized beta weights and p-values for negative binomial regression models of Y2 SOR by individual risk variable. Statistically significant effects are shown in bold.*

| | **Bivariate Model** | | | **Y2 age included as covariate** | | | **Sex included as covariate** | | |
|---|---|---|---|---|---|---|---|---|---|
| **Predictor** | **Beta Weight** | **P-Value** | **N** | **Beta Weight** | **P-Value** | **N** | **Beta Weight** | **P-Value** | **N** |
| **Prenatal Social Disadvantage** | **0.32** | **0.007** | 266 | **0.33** | **0.03** | 161 | **0.32** | **0.006** | 266 |
| **Prenatal Maternal Psychosocial Stress** | **0.26** | **0.008** | 266 | **0.29** | **0.04** | 161 | **0.26** | **0.008** | 266 |
| Birthweight | -0.0003 | 0.15 | 266 | - | - | - | - | - | - |
| Gestational Age at Birth | 0.001 | 0.99 | 266 | - | - | - | - | - | - |

| | | | | | | | | | |
|---|---|---|---|---|---|---|---|---|---|
| Sex | -0.01 | 0.96 | 266 | - | - | - | - | - | - |
| **Sensory Over-Responsivity at One Year of Age** | **1.52** | **0.0001** | 223 | **1.60** | **0.004** | 128 | **1.54** | **0.00009** | 223 |

**Table S7. SOR at age two relates to concurrent behavioral problems after accounting for covariates.** *Multivariate analyses of Y2 SOR relationship with Y2 CBCL problem scales after inclusion of key covariates (SD, age at year two, and sex included as covariates in independent models). Unstandardized beta weights and p-values are reported for negative binomial regressions between Y2 SOR and each problem scale score. Rows in bold denote significant relationships.*

| | SD included as covariate | | | Y2 age included as covariate | | | Sex included as covariate | | |
|---|---|---|---|---|---|---|---|---|---|
| **Outcome Variable** | **Beta Weight** | **P-value** | **N** | **Beta Weight** | **P-value** | **N** | **Beta Weight** | **P-value** | **N** |
| **Y2 Internalizing Problems** | **1.04** | **0.000004** | 141 | **1.05** | **0.000001** | 141 | **1.07** | **0.0000003** | 141 |
| **Y2 Externalizing Problems** | **0.60** | **0.01** | 141 | **0.53** | **0.02** | 141 | **0.53** | **0.02** | 141 |
| **Y2 Affective Problems** | **0.79** | **0.03** | 141 | **0.78** | **0.02** | 141 | **0.80** | **0.02** | 141 |
| **Y2 Anxiety Problems** | **0.79** | **0.0003** | 141 | **0.93** | **0.000006** | 141 | **0.95** | **0.000006** | 141 |
| **Y2 Autism Spectrum Problems** | **1.04** | **0.0003** | 141 | **0.92** | **0.0007** | 141 | **0.92** | **0.0007** | 141 |
| **Y2 ADHD Problems** | **0.51** | **0.01** | 141 | **0.44** | **0.02** | 141 | **0.46** | **0.02** | 141 |
| Y2 ODD Problems | - | - | - | - | - | - | - | - | - |

**Table S8. SOR at age two relates to behavioral problems when accounting for SD in inclusive sample.** *Bivariate and multivariate analyses of Y2 SOR relationship with Y2 CBCL problem scales using sample of infants born at or above 30 weeks gestational age (prenatal social disadvantage included as a covariate in independent model). Unstandardized beta weights and p-values are reported for negative binomial regressions between Y2 SOR and each problem scale score. Rows in bold denote significant relationships.*

| | Bivariate Model | SD included as covariate |
|---|---|---|

| Outcome Variable | Beta Weight | P-value | N | Beta Weight | P-value | N |
|---|---|---|---|---|---|---|
| **Y2 Internalizing Problems** | **0.92** | **0.000005** | 160 | **0.88** | **0.00004** | 160 |
| **Y2 Externalizing Problems** | **0.41** | **0.048** | 160 | **0.48** | **0.03** | 160 |
| **Y2 Affective Problems** | **0.70** | **0.03** | 160 | **0.69** | **0.04** | 160 |
| **Y2 Anxiety Problems** | **0.87** | **0.00001** | 160 | **0.70** | **0.0006** | 160 |
| **Y2 Autism Spectrum Problems** | **0.65** | **0.01** | 160 | **0.73** | **0.006** | 160 |
| Y2 ADHD Problems | 0.32 | 0.06 | 160 | - | - | - |
| Y2 ODD Problems | 0.15 | 0.57 | 160 | - | - | - |

**Table S9. SOR at age two relates to behavioral problems when accounting for age and sex in inclusive sample.** *Multivariate analyses of Y2 SOR relationship with Y2 CBCL problem scales using sample of infants born at or above 30 weeks gestational age (Y2 age and sex included as covariates in independent models). Unstandardized beta weights and p-values are reported for negative binomial regressions between Y2 SOR and each problem scale score. Rows in bold denote significant relationships.*

| | Y2 age included as covariate | | | Sex included as covariate | | |
|---|---|---|---|---|---|---|
| **Outcome Variable** | **Beta Weight** | **P-value** | **N** | **Beta Weight** | **P-value** | **N** |
| **Y2 Internalizing Problems** | **0.91** | **0.000007** | 160 | **0.93** | **0.000002** | 160 |
| **Y2 Externalizing Problems** | **0.41** | **0.046** | 160 | **0.42** | **0.04** | 160 |
| **Y2 Affective Problems** | **0.69** | **0.02** | 160 | **0.74** | **0.01** | 160 |
| **Y2 Anxiety Problems** | **0.84** | **0.000009** | 160 | **0.87** | **0.000005** | 160 |
| **Y2 Autism Spectrum Problems** | **0.64** | **0.01** | 160 | **0.65** | **0.01** | 160 |

| | | | | | | |
|---|---|---|---|---|---|---|
| Y2 ADHD Problems | - | - | - | - | - | - |
| Y2 ODD Problems | - | - | - | - | - | - |

**Table S10. SOR at age two predicts behavioral problems after accounting for SD and sex.** *Multivariate analyses of Y2 SOR relationship with Y3 CBCL problem scales after inclusion of key covariates (SD and sex included as covariates in independent models). Unstandardized beta weights and p-values are reported for negative binomial regressions between Y2 SOR and each problem scale score. Rows in bold denote significant relationships.*

| | SD included as covariate | | | Sex included as covariate | | |
|---|---|---|---|---|---|---|
| **Outcome Variable** | **Beta Weight** | **P-value** | **N** | **Beta Weight** | **P-value** | **N** |
| **Y3 Internalizing Problems** | **0.84** | **0.000006** | 164 | **0.85** | **0.000001** | 164 |
| **Y3 Externalizing Problems** | **0.38** | **0.04** | 164 | **0.41** | **0.02** | 164 |
| **Y3 Affective Problems** | **0.52** | **0.03** | 164 | **0.56** | **0.02** | 164 |
| **Y3 Anxiety Problems** | **0.75** | **0.0005** | 164 | **0.78** | **0.0001** | 164 |
| **Y3 Autism Spectrum Problems** | **0.95** | **0.00001** | 164 | **0.94** | **0.000003** | 164 |
| **Y3 ADHD Problems** | 0.28 | 0.08 | 164 | **0.37** | **0.01** | 164 |
| Y3 ODD Problems | - | - | - | - | - | - |

**Table S11. SOR at age two predicts behavioral problems after accounting for age.** *Multivariate analyses of Y2 SOR relationship with Y3 CBCL problem scales after inclusion of key covariates (Y2 age and Y3 age included as covariates in independent models). Unstandardized beta weights and p-values are reported for negative binomial regressions between Y2 SOR and each problem scale score. Rows in bold denote significant relationships.*

| | Y2 age included as covariate | | | Y3 age included as covariate | | |
|---|---|---|---|---|---|---|
| **Outcome Variable** | **Beta Weight** | **P-value** | **N** | **Beta Weight** | **P-value** | **N** |
| **Y3 Internalizing Problems** | **1.18** | **0.000002** | 97 | **0.84** | **0.000002** | 164 |
| **Y3 Externalizing Problems** | **0.59** | **0.02** | 97 | **0.36** | **0.03** | 164 |
| **Y3 Affective Problems** | **0.84** | **0.003** | 97 | **0.53** | **0.02** | 164 |
| **Y3 Anxiety Problems** | **0.95** | **0.0004** | 97 | **0.78** | **0.0001** | 164 |
| **Y3 Autism Spectrum Problems** | **1.15** | **0.00009** | 97 | **0.92** | **0.000006** | 164 |
| **Y3 ADHD Problems** | **0.51** | **0.02** | 97 | **0.34** | **0.02** | 164 |
| Y3 ODD Problems | - | - | - | - | - | - |

**Table S12. SOR at age two predicts behavioral problems in inclusive sample after accounting for previous symptoms and SD.** *Bivariate and multivariate analyses of Y2 SOR relationship with Y3 CBCL problem scales using a sample of infants born at or above 30 weeks gestational age (Y2 symptoms and prenatal social disadvantage included as covariates in independent models). Unstandardized beta weights and p-values are reported for negative binomial regressions between Y2 SOR and each problem scale score. Rows in bold denote significant relationships.*

| | Bivariate Model | | | Y2 Sx included as covariate | | | SD included as covariate | | |
|---|---|---|---|---|---|---|---|---|---|
| **Outcome Variable** | **Beta Weight** | **P-value** | **N** | **Beta Weight** | **P-value** | **N** | **Beta Weight** | **P-value** | **N** |
| **Y3 Internalizing Problems** | **0.74** | **0.00001** | 186 | **0.60** | **0.007** | 110 | **0.70** | **0.00009** | 186 |
| Y3 Externalizing Problems | 0.26 | 0.11 | 186 | - | - | - | - | - | - |
| Y3 Affective Problems | 0.42 | 0.052 | 186 | - | - | - | - | - | - |
| **Y3 Anxiety Problems** | **0.74** | **0.00007** | 186 | **0.46** | **0.046** | 110 | **0.64** | **0.001** | 186 |
| **Y3 Autism Spectrum Problems** | **0.76** | **0.0001** | 186 | **0.61** | **0.02** | 110 | **0.76** | **0.0003** | 186 |
| Y3 ADHD Problems | 0.24 | 0.08 | 186 | - | - | - | - | - | - |
| Y3 ODD Problems | 0.14 | 0.47 | 186 | - | - | - | - | - | - |

**Table S13. SOR at age two predicts behavioral problems in inclusive sample after accounting for sex and age.** *Multivariate analyses of Y2 SOR relationship with Y3 CBCL problem scales in a sample of infants born at or above 30 weeks gestational age (sex, Y2 age, and Y3 age included as covariates in independent models). Unstandardized beta weights and p-values are reported for negative binomial regressions between Y2 SOR and each problem scale score. Rows in bold denote significant relationships.*

| | Sex included as covariate | | | Y2 age included as covariate | | | Y3 age included as covariate | | |
|---|---|---|---|---|---|---|---|---|---|
| **Outcome Variable** | **Beta Weight** | **P-value** | **N** | **Beta Weight** | **P-value** | **N** | **Beta Weight** | **P-value** | **N** |
| **Y3 Internalizing Problems** | **0.73** | **0.00001** | 186 | **0.92** | **0.0001** | 111 | **0.73** | **0.00002** | 186 |
| Y3 Externalizing Problems | - | - | - | - | - | - | - | - | - |

| | | | | | | | | | |
|---|---|---|---|---|---|---|---|---|---|
| Y3 Affective Problems | - | - | - | - | - | - | - | - | - |
| **Y3 Anxiety Problems** | **0.73** | **0.00009** | 186 | **0.83** | **0.0007** | 111 | **0.71** | **0.0002** | 186 |
| **Y3 Autism Spectrum Problems** | **0.76** | **0.0001** | 186 | **0.78** | **0.006** | 111 | **0.75** | **0.0002** | 186 |
| Y3 ADHD Problems | - | - | - | - | - | - | - | - | - |
| Y3 ODD Problems | - | - | - | - | - | - | - | - | - |

**Table S14. SOR at age two predicts greater internalizing symptoms when accounting for age.** *Multivariate analyses of Y2 SOR relationship with preschool-age (ages 4-6) PAPA symptoms (age at PAPA administration included as a covariate in an independent model). Unstandardized beta weights and p-values are reported for negative binomial regressions between Y2 SOR and each symptom number. Rows in bold denote significant relationships.*

| | **PAPA age included as covariate** | | |
|---|---|---|---|
| **Outcome Variable** | **Beta Weight** | **P-value** | **N** |
| **Number of Internalizing Symptoms** | **0.40** | **0.02** | 180 |
| Number of Externalizing Symptoms | - | - | - |
| **Number of Separation Anxiety Symptoms** | **0.72** | **0.008** | 180 |
| **Number of Social Phobia Symptoms** | **1.38** | **0.03** | 180 |
| **Number of Generalized Anxiety Symptoms** | **0.38** | **0.03** | 180 |

**Table S15. SOR at age two predicts greater separation anxiety symptoms in inclusive sample.** *Bivariate and multivariate analyses of Y2 SOR relationship with preschool-age (ages 4-6) PAPA symptoms in a sample of infants born at or above 30 weeks gestational age. Unstandardized beta weights and p-values are reported for negative binomial regressions between Y2 SOR and each symptom number. Rows in bold denote significant relationships.*

| | **Bivariate Model** | | | **PAPA age included as covariate** | | |
|---|---|---|---|---|---|---|
| **Outcome Variable** | **Beta Weight** | **P-value** | **N** | **Beta Weight** | **P-value** | **N** |
| Number of Internalizing Symptoms | 0.25 | 0.14 | 207 | - | - | - |
| Number of Externalizing Symptoms | 0.18 | 0.49 | 207 | - | - | - |
| **Number of Separation Anxiety Symptoms** | **0.54** | **0.04** | 207 | **0.55** | **0.04** | 207 |
| Number of Social Phobia Symptoms | 0.93 | 0.12 | 207 | - | - | - |
| Number of Generalized Anxiety Symptoms | 0.22 | 0.18 | 207 | - | - | - |

**Table S16. SOR at age two predicts anxiety diagnoses after accounting for age.** *Multivariate analyses of Y2 SOR relationship with preschool-age PAPA diagnoses (age at PAPA administration included as a covariate in an independent model). Unstandardized beta weights and p-values are reported for logistic regressions between Y2 SOR and each diagnosis variable. Rows in bold denote significant relationships.*

| | **PAPA age included as covariate** | | |
|---|---|---|---|
| **Outcome Diagnosis** | **Beta Weight** | **P-value** | **N** |
| Any Internalizing Disorder | - | - | - |
| Any Externalizing Disorder | - | - | - |
| **Separation Anxiety** | **1.44** | **0.03** | 180 |
| **Social Phobia** | **1.42** | **0.04** | 180 |
| Generalized Anxiety Disorder | - | - | - |

**Table S17. SOR at age two does not predict diagnoses in inclusive sample.** *Bivariate analyses of Y2 SOR relationship with preschool-age PAPA diagnoses in a sample of children born at or above 30 weeks gestational age. Unstandardized beta weights and p-values are reported for logistic regressions between Y2 SOR and each diagnosis variable. No relationships were found to be significant.*

| **Outcome Diagnosis** | **Beta Weight** | **P-value** | **N** |
|---|---|---|---|
| Any Internalizing Disorder | 0.30 | 0.53 | 207 |
| Any Externalizing Disorder | -0.71 | 0.23 | 207 |
| Separation Anxiety | 1.03 | 0.09 | 207 |
| Social Phobia | 1.01 | 0.10 | 207 |
| Generalized Anxiety Disorder | 0.19 | 0.69 | 207 |

**Table S18. Relations between FC and infant-toddler SOR after accounting for key covariates.** *Multivariate analyses of infant-toddler SOR relationship with neonatal functional connectivity (frames retained, prenatal social disadvantage, and prenatal psychosocial stress included as covariates in independent models). Unstandardized beta weights and p-values are reported for general linear regressions between SOR and functional connectivity pair. Bold numbers denote significant relationships.*

| | **Frames Retained included as covariate** | | | **SD included as covariate** | | | **PSS included as covariate** | | |
|---|---|---|---|---|---|---|---|---|---|
| **Outcome Variable** | **Beta Weight** | **P-value** | **N** | **Beta Weight** | **P-value** | **N** | **Beta Weight** | **P-value** | **N** |
| **Thalamus vs. Auditory Cortex** | **0.04** | **0.047** | 236 | 0.03 | 0.16 | 236 | 0.03 | 0.15 | 236 |

| | | | | | | | | | |
|---|---|---|---|---|---|---|---|---|---|
| **Thalamus vs. Somatomotor Hand** | **0.04** | **0.01** | 236 | **0.03** | **0.048** | 236 | **0.04** | **0.02** | 236 |
| Caudate vs. Auditory Cortex | - | - | - | - | - | - | - | - | - |
| Caudate vs. Cinguloparietal | - | - | - | - | - | - | - | - | - |
| Caudate vs. Somatomotor Hand | - | - | - | - | - | - | - | - | - |
| Somatomotor Hand vs. Somatomotor Hand | - | - | - | - | - | - | - | - | - |
| Somatomotor Mouth vs. Somatomotor Hand | - | - | - | - | - | - | - | - | - |
| Salience vs. Somatomotor Hand | - | - | - | - | - | - | - | - | - |
| Ventral Attention Network vs. Ventral Attention Network | - | - | - | - | - | - | - | - | - |

**Table S19. Relations between neonatal FC and infant-toddler SOR in inclusive sample.** *Bivariate analyses of composite SOR relationship with network-level FC pairs in a sample of infants born at or above 30 weeks gestational age. Unstandardized beta weights and p-values are reported for general linear regressions between SOR and each functional connectivity pair. A strong, positive relationship is observed between SOR and Thalamus vs. Somatomotor Hand in a bivariate model. Rows in bold denote significant relationships.*

| Outcome FC Pair | Beta Weight | P-value | N |
|---|---|---|---|
| Caudate vs. Cinguloparietal | 0.006 | 0.67 | 275 |
| **Thalamus vs. Auditory Cortex** | **0.04** | **0.03** | 275 |
| Caudate vs. Auditory Cortex | 0.01 | 0.42 | 275 |
| **Thalamus vs. Somatomotor Hand** | **0.03** | **0.02** | 275 |
| Caudate vs. Somatomotor Hand | -0.0002 | 0.98 | 275 |
| Somatomotor Hand vs. Somatomotor Hand | 0.008 | 0.41 | 275 |
| Somatomotor Mouth vs. Somatomotor Hand | 0.02 | 0.21 | 275 |
| Salience vs. Somatomotor Hand | -0.003 | 0.74 | 275 |
| Ventral Attention Network vs. Ventral Attention Network | -0.01 | 0.23 | 275 |

**Table S20. Relations between neonatal FC and infant-toddler SOR in inclusive sample after accounting for key covariates.** *Multivariate analyses of infant-toddler SOR relationship with neonatal functional connectivity in a sample of infants born at or above 30 weeks gestational age (frames retained, prenatal social disadvantage, and prenatal psychosocial stress included as covariates in independent models). Unstandardized beta weights and p-values are reported for general linear regressions between SOR and each functional connectivity pair. Bold numbers denote significant relationships.*

| | Frames Retained included as covariate | | | SD included as covariate | | | PSS included as covariate | | |
|---|---|---|---|---|---|---|---|---|---|
| **Outcome Variable** | **Beta Weight** | **P-value** | **N** | **Beta Weight** | **P-value** | **N** | **Beta Weight** | **P-value** | **N** |
| **Thalamus vs. Auditory Cortex** | **0.03** | **0.04** | 275 | 0.03 | 0.12 | 275 | 0.03 | 0.08 | 275 |
| **Thalamus vs. Somatomotor Hand** | **0.03** | **0.02** | 275 | 0.02 | 0.09 | 275 | **0.03** | **0.03** | 275 |
| Caudate vs. Auditory Cortex | - | - | - | - | - | - | - | - | - |
| Caudate vs. Cinguloparietal | - | - | - | - | - | - | - | - | - |
| Caudate vs. Somatomotor Hand | - | - | - | - | - | - | - | - | - |
| Somatomotor Hand vs. Somatomotor Hand | - | - | - | - | - | - | - | - | - |
| Somatomotor Mouth vs. Somatomotor Hand | - | - | - | - | - | - | - | - | - |
| Salience vs. Somatomotor Hand | - | - | - | - | - | - | - | - | - |
| Ventral Attention Network vs. Ventral Attention Network | - | - | - | - | - | - | - | - | - |

**Supplemental Figures**

**Figure S1. Gestational age in the full sample.** (A) *Histogram depicting the frequency of observed gestational age (in weeks, unwinsorized) in the full sample. (B) Scatterplot depicting the relationship between gestational age (in weeks) and Y2 SOR.*

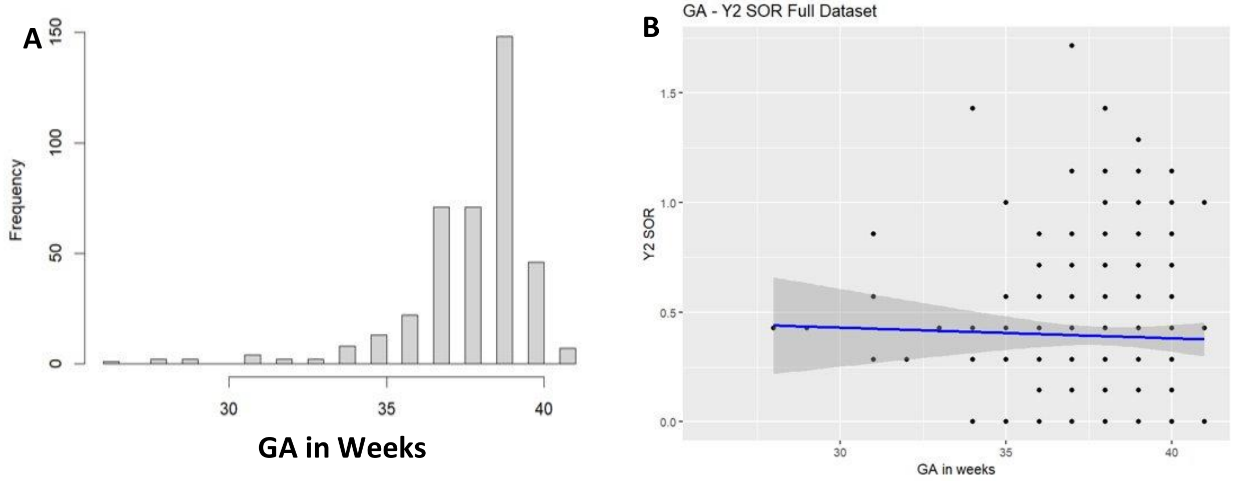


**Figure S2. Spearman's correlations between predictor variables.** *Spearman's correlation coefficients are displayed to indicate the relationship between each predictor variable. Blue indicates positively correlated measures, and red indicates negatively correlated measures.*

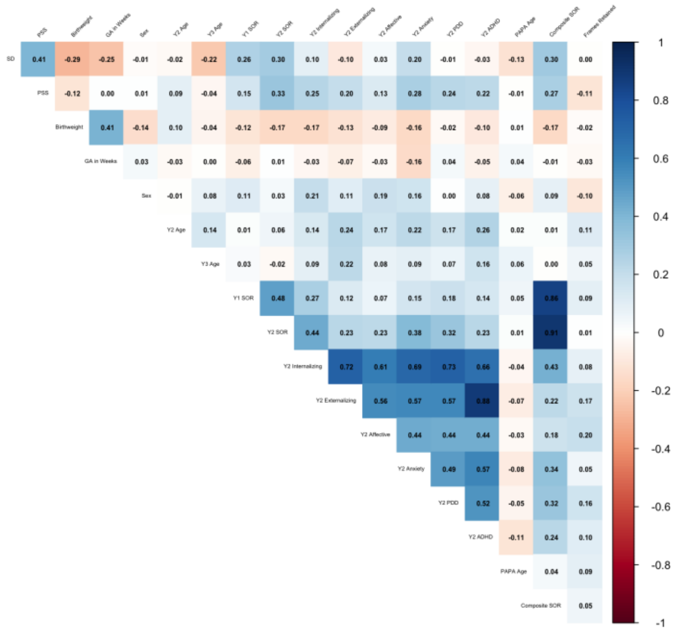


**Figure S3. SOR distribution at years one and two.** *The x-axis represents mean ITSEA sensory sensitivity subscale score (winsorized), and each ridge shows the distribution of scores at a given year. The plot highlights the left skew across timepoints, while also displaying a greater range of*

*values, particularly more participants reporting greater SOR, in year two scores as opposed to year one scores.*

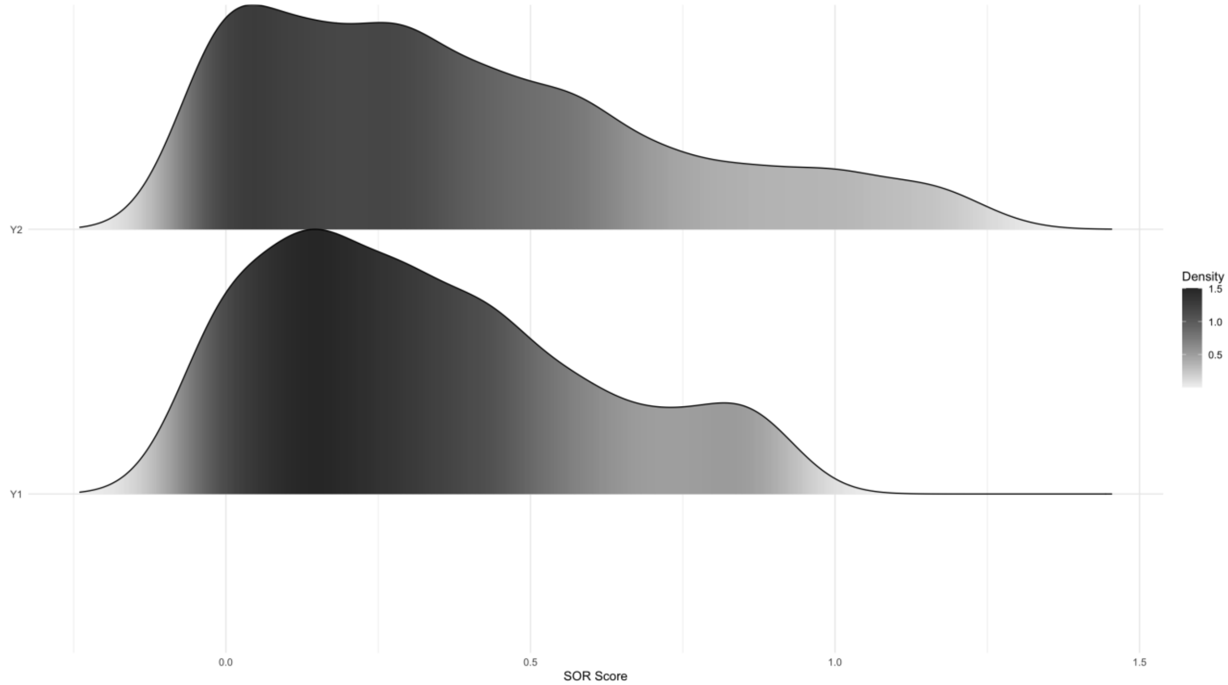


**Figure S4. Internalizing and externalizing score distributions at years two and three.** *The x-axes represent winsorized CBCL internalizing and externalizing domain scores, and each ridge shows the distribution at a given timepoint. (A) The upper plot displays a left-skew of internalizing domain scores across years two and three, with the range of scores being greater at year three as compared to year two. (B) The lower plot highlights a left-skew of externalizing domain scores across years two and three, with the range of scores increased at year three compared to year one.*

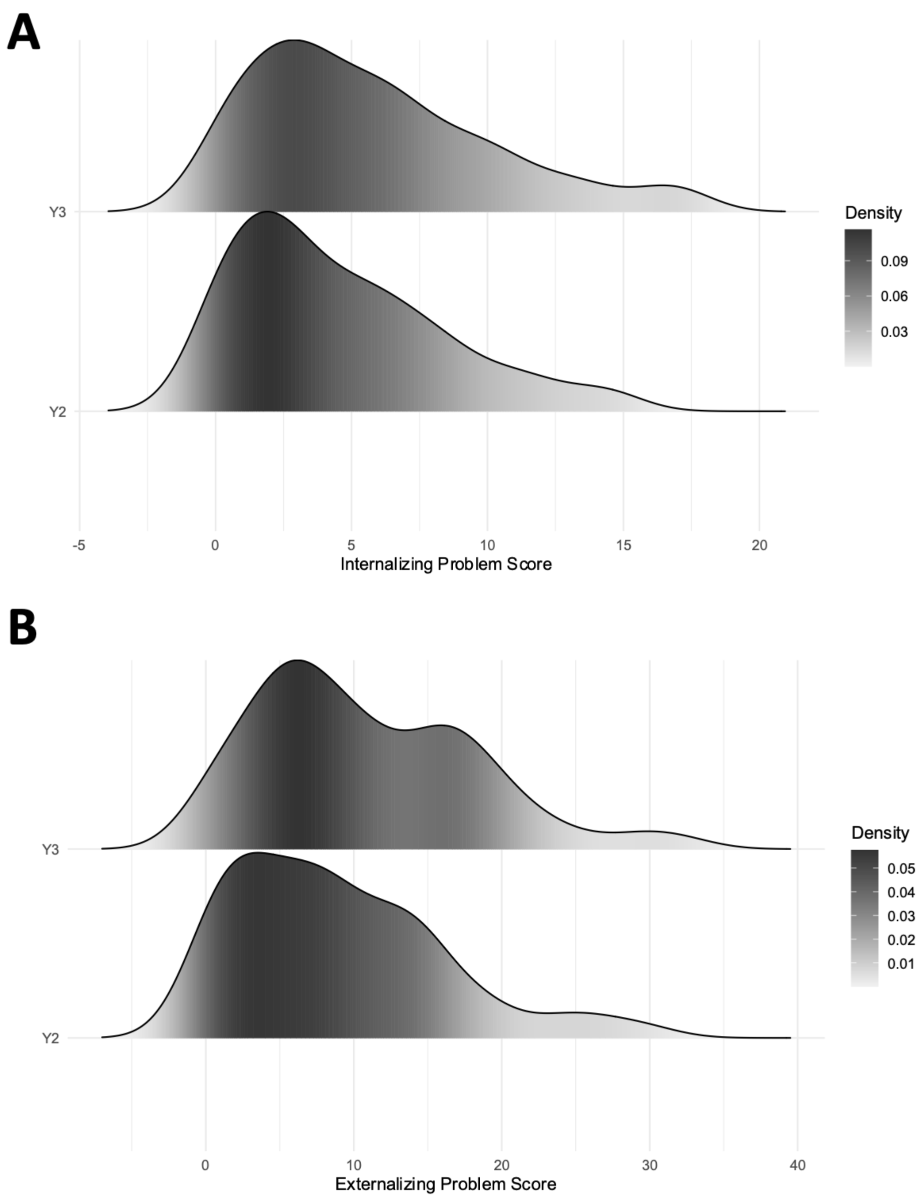


**Figure S5. Y2 SOR mediates associations of prenatal adversity with Y3 behavioral problem scales.** *(a): Path diagram of mediation model. Prenatal psychosocial stress (IV) predicts Y3 internalizing problems (DV) both directly and indirectly through Y2 SOR (mediator). Unstandardized coefficients are displayed on each path, with significance levels indicated. The indirect path through Y2 SOR was significant, suggesting partial mediation by Y2 SOR. (b): Path diagram of mediation model. Prenatal psychosocial stress (IV) predicts Y3 externalizing problems (DV) directly but not indirectly through Y2 SOR (mediator). Unstandardized coefficients are displayed on each path, with significance levels indicated. The indirect path through Y2 SOR was not significant, suggesting no mediating effect of Y2 SOR on the relationship between psychosocial stress and Y3 externalizing problems.*

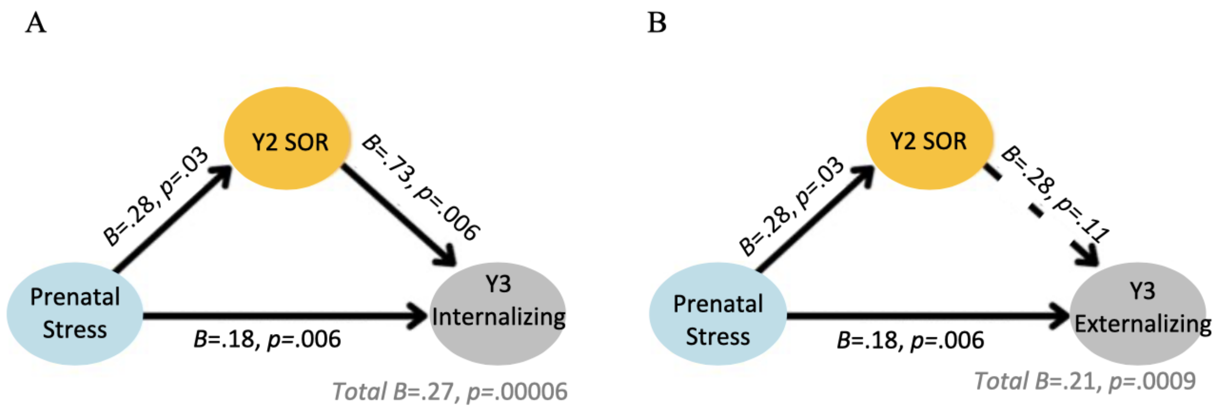


**Figure S6. Y2 SOR mediates associations of prenatal adversity with Y3 DSM-oriented scales in inclusive sample.** *Mediation analyses performed using a sample of infants born at or above 30 weeks gestational age. (a): Path diagram of mediation model. Prenatal psychosocial stress (IV) predicts Y3 anxiety problems (DV) both directly and indirectly through Y2 SOR (mediator). Unstandardized coefficients are displayed on each path, with significance levels indicated. The indirect path through Y2 SOR was significant, indicating partial mediation by Y2 SOR. (b) Path diagram of mediation model. Prenatal psychosocial stress (IV) predicts Y3 autism-spectrum problems (DV) both directly and indirectly through Y2 SOR (mediator). Unstandardized coefficients are displayed on each path, with significance levels indicated. The indirect path through Y2 SOR was significant, indicating partial mediation by Y2 SOR. (c). Path diagram of mediation model. Prenatal social disadvantage (IV) predicts Y3 anxiety problems (DV) indirectly through Y2 SOR (mediator). Unstandardized coefficients are displayed on each path with significance levels indicated. The total indirect effect was significant, suggesting full mediation by Y2 SOR.*

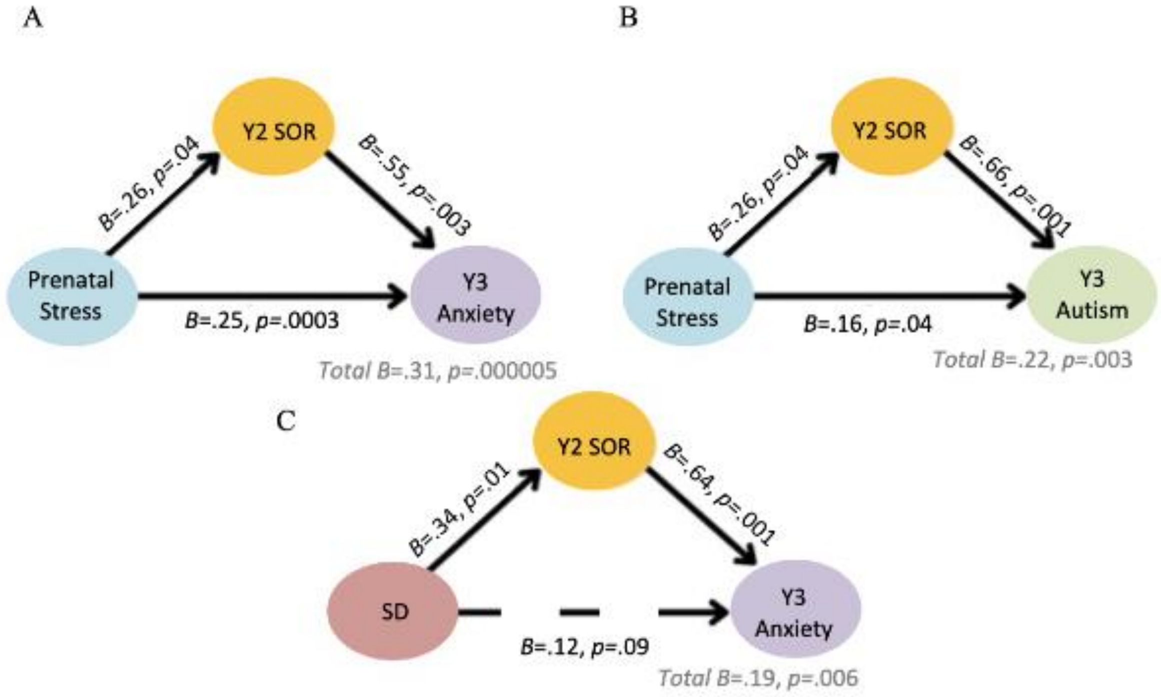


**Figure S7. Y2 SOR mediates associations of prenatal adversity with Y3 behavioral problem scales in inclusive sample.** *Mediation analyses performed using a sample of infants born at or above 30 weeks gestational age. (a): Path diagram of mediation model. Prenatal psychosocial stress (IV) predicts Y3 internalizing problems (DV) both directly and indirectly through Y2 SOR (mediator). Unstandardized coefficients are displayed on each path, with significance levels indicated. The indirect path through Y2 SOR was significant, suggesting partial mediation by Y2 SOR. (b): Path diagram of mediation model. Prenatal psychosocial stress (IV) predicts Y3 externalizing problems (DV) directly but not indirectly through Y2 SOR (mediator). Unstandardized coefficients are displayed on each path, with significance levels indicated. The indirect path through Y2 SOR was not significant, suggesting no mediating effect of Y2 SOR on the relationship between psychosocial stress and Y3 externalizing problems.*

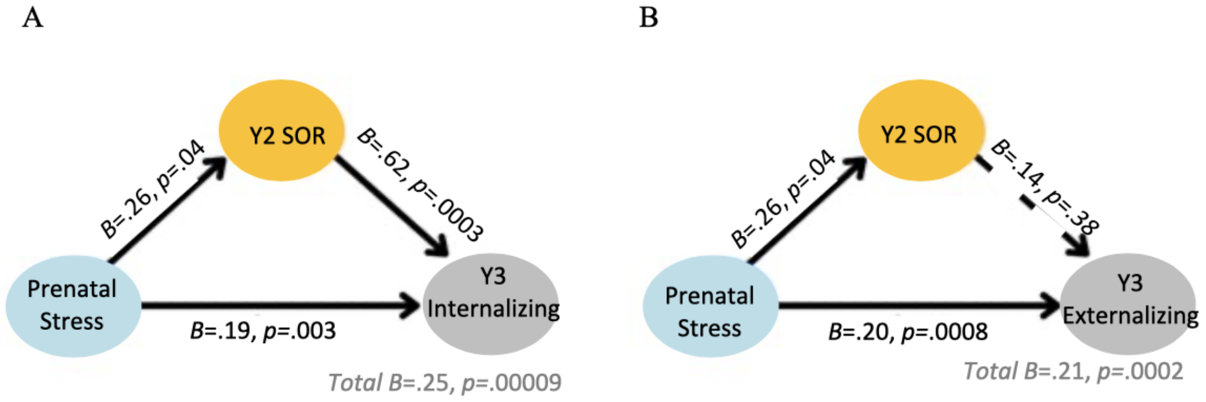

### Supplemental References


Achenbach, T. M., & Rescorla, L. A. (2000). *Manual for the ASEBA preschool forms and profiles* (Vol. 30). Burlington, VT: University of Vermont, Research center for children, youth ….

Carter, A. S., Briggs-Gowan, M. J., Jones, S. M., & Little, T. D. (2003). The Infant-Toddler Social and Emotional Assessment (ITSEA): Factor Structure, Reliability, and Validity. *Journal of Abnormal Child Psychology*, *31*(5), 495–514.

Carter, A. S., Little, C., Briggs-Gowan, M. J., & Kogan, N. (1999). The infant-toddler social and emotional assessment (ITSEA): Comparing parent ratings to laboratory observations of task mastery, emotion regulation, coping behaviors, and attachment status. *Infant Mental Health Journal*, *20*(4), 375–392.

Cohen, S., Kamarck, T., & Mermelstein, R. (1983). A global measure of perceived stress. *Journal of Health and Social Behavior*, *24*, 386–396.

Cox, J. L., Holden, J. M., & Sagovsky, R. (1987). Detection of Postnatal Depression: Development of the 10-item Edinburgh Postnatal Depression Scale. *British Journal of Psychiatry*, *150*(6), 782–786. Cambridge Core. https://doi.org/10.1192/bjp.150.6.782

Egger, H. L., Angold, A., Small, B., & Copeland, W. (1999). The preschool age psychiatric assessment. In *The Oxford Handbook of Infant, Toddler, and Preschool Mental Health Assessment*.

Guenther, P. M., Kirkpatrick, S. I., Reedy, J., Krebs-Smith, S. M., Buckman, D. W., Dodd, K. W., Casavale, K. O., & Carroll, R. J. (2014). The Healthy Eating Index-2010 is a valid and reliable measure of diet quality according to the 2010 Dietary Guidelines for Americans. *The Journal of Nutrition*, *144*(3), 399–407.

Helen Egger L., Angold, A., Small, B., & Copeland, W. (2020). Preschool Age Psychiatric Assessment: A Structured Parent Interview for Assessing Psychiatric Symptoms and Disorders in Preschool Children. In *The Oxford Handbook of Infant, Toddler, and Preschool Mental Health Assessment, 2nd Edition* (2nd ed.). Oxford University Press.

Kind, A. J., & Buckingham, W. R. (2018). Making neighborhood-disadvantage metrics accessible—The neighborhood atlas. *The New England Journal of Medicine*, *378*(26), 2456.

Lewis, T. T., Yang, F. M., Jacobs, E. A., & Fitchett, G. (2012). Racial/ethnic differences in responses to the everyday discrimination scale: A differential item functioning analysis. *American Journal of Epidemiology*, *175*(5), 391–401.

Slavich, G. M., & Shields, G. S. (2018). Assessing Lifetime Stress Exposure Using the Stress and Adversity Inventory for Adults (Adult STRAIN): An Overview and Initial Validation. *Psychosomatic Medicine*, *80*(1), 17–27. https://doi.org/10.1097/PSY.0000000000000534
